# A comparative GWAS of eye colour in light and dark eye genetic backgrounds defined by *HERC2* rs12913832 polymorphism

**DOI:** 10.1101/2025.07.20.665796

**Authors:** Cristina L Abbatangelo, Frida Lona Durazo, Melissa Edwards, Esteban J Parra

**Affiliations:** Department of Anthropology, Faculty of Arts and Science, University of Toronto, Toronto, ON, Canada; Montréal Heart Institute, Montréal, QC, Canada; Université de Montréal, Montréal, QC, Canada; Brain Tumour Research Center, SickKids Research Institute, Peter Gilgan Center for Research and Learning, Toronto, ON, Canada; Department of Anthropology, University of Toronto Mississauga, Mississauga, ON, Canada

**Keywords:** rs12913832, eye colour, genome-wide association study (GWAS), *HERC2*, *OCA2*, quantitative eye colour measures, CanPath

## Abstract

rs12913832, a polymorphism located in an enhancer within the *HERC2* gene, which is known to regulate *OCA2* transcription, is heavily relied upon as a predictor of light versus dark eyes. Individuals with the GG genotype are projected to have blue eyes, while individuals with the AA or AG genotypes are projected to have darker eye colours (primarily brown). However, eye colour is a polygenic trait, and previous studies have revealed that a significant proportion of individuals self-report an eye colour that is not concordant with their genotype at rs12913832. Herein, we address the question: What common markers are influencing eye colour in individual’s whose self-reported phenotype does not correspond to the expected phenotype based on their rs12913832 genotype? Building upon our prior investigation of iris pigmentation genetics in individuals with an expected “blue eye” background (rs12913832:GG genotype) in a sample of the Canadian Partnership for Tomorrow’s Health (CanPath) cohort, this study extends the analysis to include individuals with an expected “brown eye” background (rs12913832:AA+AG). We identified variants in *SLC45A2*, *TYRP1*, *TYR*, *SLC24A4* and *TSPAN10*, which may influence eye colour presentation in individuals with the rs12913832:GG genotype and variants in *IRF4*, *TYRP1* and *OCA2*, which may influence eye colour presentation in individuals with the rs12913832:AA+AG genotype. These markers include well-known pigmentation-associated single nucleotide polymorphisms, such as rs16891982 (*SLC45A2*), rs1126809 (*TYR*), rs12203592 (*IRF4*), rs1800407 (*OCA2*) and rs6420484 (*TSPAN10*). Several of these loci were replicated using independent quantitative eye colour measures, including heterochromia and CIELAB colour dimensions. This research highlights the importance of gene-gene interactions and the polygenic nature of pigmentation traits, emphasizing modifying effects that can sometimes counteract the dominant influence of rs12913832, contributing to advancements in pigmentation genetics and forensic applications.

## Introduction

Human pigmentation traits, such as hair, skin, and eye colour, represent some of the most striking examples of visible genetic variation. These traits are highly polygenic, with complex gene-gene and gene-environment interactions contributing to their phenotypic expression. Understanding the genetic architecture underlying these traits not only informs biological mechanisms but also enhances applications such as predictive modeling, reconstruction of ancestral phenotypes and human identification. Of particular interest is the genetic basis underlying iris pigmentation, which remains incompletely understood despite significant advances in large-scale genome-wide association studies (GWAS) (1).

A key genetic determinant of human eye colour is the single nucleotide polymorphism (SNP) rs12913832, located in a conserved enhancer region within the *HERC2* gene that regulates transcription of the neighbouring *OCA2* gene (2–5). *OCA2* plays a central role in melanin biosynthesis, and its expression levels are strongly influenced by allelic variation at rs12913832. The ancestral A allele facilitates transcription factor binding and promotes long-range chromatin looping, enabling physical interaction between the *OCA2* promoter and the *HERC2* enhancer. This configuration enhances *OCA2* expression, leading to increased melanin production and resulting in darker eye colours, such as brown and hazel (4,6). Conversely, the derived G allele reduces chromatin looping efficiency, thereby decreasing *OCA2* expression and melanin production. This reduction is associated with lighter iris pigmentation, characteristic of blue eyes (4,6). As such, rs12913832 has become a key marker in pigmentation research and forensic DNA phenotyping (FDP), due to its strong predictive value for distinguishing between light and dark eye colours (2,4,7). Individuals with the GG genotype are typically predicted to have blue eyes, while those with AG or AA genotypes are generally expected to exhibit brown eyes (**Fig 1**). However, while these patterns hold true for most individuals, particularly within European populations where eye colour diversity is greatest, genotype-phenotype mismatches have been reported.

**Fig 1.**
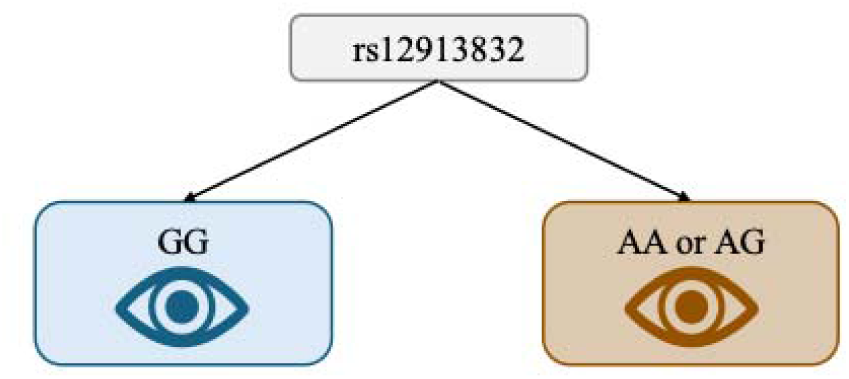
Expected eye colour phenotype based on rs12913832 genotype.

Specifically, previous research has illustrated that some rs12913832:GG individuals present with eye colours other than blue, and some rs12913832:AA or AG individuals show eye colours other than brown (8–10). A recent publication found a notable discrepancy in the Canadian Partnership for Tomorrow’s Health (CanPath) cohort where 33% of individuals with the rs12913832:GG genotype self-reported eye colours other than blue (11). This finding challenges the notion that the rs12913832:GG genotype exclusively corresponds to blue eyes and highlights the presence of additional genetic or regulatory factors that may modulate melanin production, resulting in a broader spectrum of iris pigmentation. In a previous study (12), we stratified participants of the CanPath cohort based on rs12913832 genotypes and identified a number of SNPs that seem to modify the effect of the rs12913832 GG genotype, contributing to an explanation as to why individuals self-report “non-blue” eye colour in spite of having severely reduced expression of the *OCA2* gene, which usually results in blue eye colour.

The present study builds on this foundation by extending the 2023 study in two important ways: 1) Identifying in the same CanPath sample used in the 2023 study potential genetic modifiers that may explain individuals with rs12913832:AA+AG genotypes reporting eye colours other than brown and hazel, and 2) Replicating the same markers in another independent sample of individuals of European ancestry, for whom detailed quantitative iris pigmentation measures are available (including measures capturing central heterochromia, defined as the presence of different eye colours in the inner and outer portions of the iris). By incorporating additional GWAS and post-GWAS analyses, this work aims to address the following research question: **What common markers are influencing eye colour variation in individuals whose self-reported phenotype does not correspond to the expected phenotype associated with their genotype at rs12913832?** Analyzing both self-reported data and detailed quantitative measures of iris colour, we describe polymorphisms that act to counteract the strong influence of rs12913832 on iris colour.

## Methods

### Study Dataset and Design

This research was conducted under the approval of the University of Toronto Research Ethics Board (REB), adhering to Human Research Protocol #36429. The study dataset was drawn from the Canadian Partnership for Tomorrow’s Health (CanPath) biobank, Canada’s largest collection of population health data, containing genomic and phenotypic information from over 300,000 Canadians of predominantly European ancestry (13,14). Access to the data was granted through CanPath, under application number DAO-034431 (Research Project Number 329058, 2018-10-01 start date); genotype and phenotype data were accessed 04/10/2021. All analyses were conducted using de-identified data. The authors did not participate in data collection and had no access to personally identifiable information at any stage of the research. All participants provided written informed consent at the time of data collection by CanPath.

The study utilized a subset of 5,481 individuals for which self-reported eye colour was available (*N_GG_*=2,724 and *N_AA+AG_*=2,757), encompassing various Canadian provinces. **Fig. 2** illustrates the provincial cohorts represented in the biobank. Among this group, 56.29% were female, and the average age was approximately 55 years (standard error ± 0.12).

**Fig 2.**
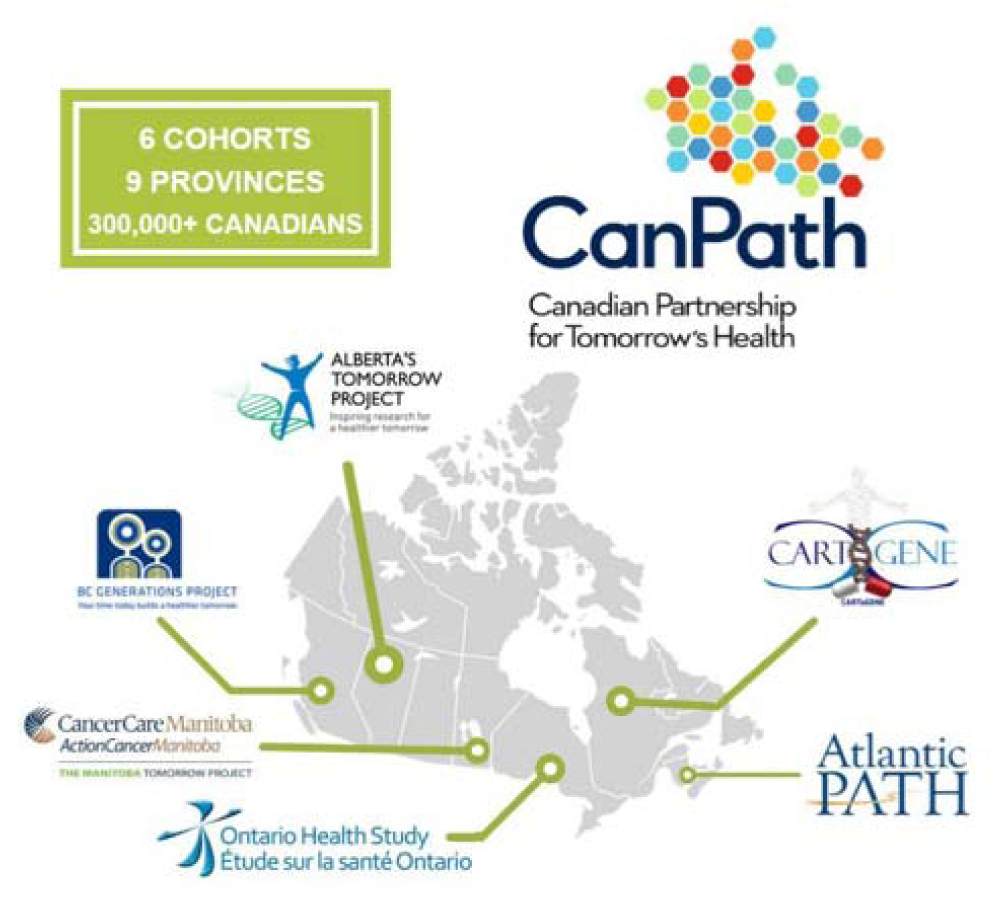
Study cohorts and provinces participating in the Canadian Partnership for Tomorrow’s Health (CanPath). Additional details about the study and participating cohorts can be found in (14) and at <u>canpath.ca</u>.

CanPath participants underwent genotyping between 2012 and 2018. Two genotyping chips were utilized: the Axiom 2.0 UK Biobank Array (Affymetrix) (UKBB) for (*N*=3,212) individuals and the Global Screening Array (GSA) 24v1+MDP (*N*=2,429) participants, conducted by CanPath. The genotyping arrays analyzed between 658,296 and 813,168 SNPs. Imputation was carried out previously on the Sanger Imputation Server using the Haplotype Reference Consortium dataset as a reference for imputation (11,12).

Among the 2,724 participants with rs12913832:AA or AG genotypes and available self-reported eye colour data, 512 reported eye colours other than brown or hazel. This observation suggests that the rs12913832:AA+AG genotype alone does not fully account for dark eye pigmentation and points to the possibility that other genetic variants may influence eye colour in individuals with the AA+AG genotype.

To investigate, we piggybacked off the 2023 rs12913832:GG study (12) and applied the same methods for AA+AG individuals in the CanPath sample, as well as a separate replication dataset. **Fig 3** illustrates the step-by-step computational methodology.

**Fig 3.**
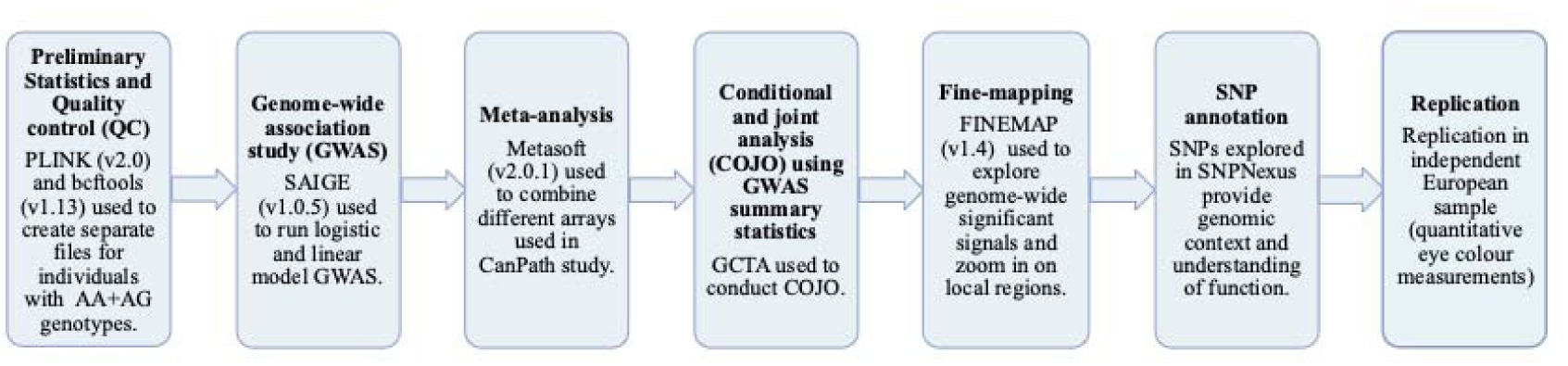
Step-by-step computational methodology. Flow chart illustrating the computational steps used to perform a genome-wide association study (GWAS) to identify genetic variants associated with eye colour in rs12913832:AA+AG individuals.

### Preliminary Statistics and Quality Control (QC)

Participants with rs12913832:AA+AG genotypes were extracted from CanPath arrays (UKBB and GSA). We retained only those individuals who self-reported as having European-related ancestry and for which self-reported eye colour data were available. Individuals who self-reported amber eyes were excluded due to the low frequency of individuals who reported this colour category.

Pre-imputation QC was performed as described in Lona-Durazo et al (2022) (11) . Post-imputation QC began with the removal of all variants other than bi-allelic SNPs (e.g., insertion/deletions, multi-allelic markers, and other types of polymorphisms) using bcftools (version 1.13) (15,16). Additionally, bcftools was also used to ensure that only SNPs were represented in the imputed genotype files, no insertion/deletions (indels) or other types of polymorphisms, as well as to remove markers with low imputation accuracy (`INFO>0.8`) and duplicated markers. Variants with strong deviations from Hardy-Weinberg equilibrium (HWE) (`--hwe 1E-6`) were filtered from the imputed files using `vcftools` (version 0.1.16.) (17). Pre-imputation and post-imputation QCs were performed separately on each array.

An initial principal component analysis (PCA) was performed using a pruned (`--indeppairwise 50 1 0.1`) set of unimputed variants from both CanPath arrays projected onto the 1000s Genomes Project (1KG) Phase 3 samples using PLINK (version 1.9) (18,19) to identify any outliers that did not cluster within the European superpopulation of the 1KGP based on the first three principal components. After removal of outliers, 1,412 samples were retained from the GSA array and 1,312 from the UKBB array (for a total of 2,724 AA/AG individuals). A second PCA was then performed for each array using PLINK (version 1.9) (18,19) and the first 10 principal components were retained as covariates and added to the phenotype files corresponding to each array using R studio (version 4.2.0) (20).

### Genome-wide Association Study (GWAS)

A genome-wide association study (GWAS) targeting eye colour was independently conducted on individuals with AA+AG genotypes from each array, employing both linear and logistic mixed models via SAIGE (version 1.0.5) (21). In the linear model, eye colour is coded as follows: blue=1, green=2, hazel=3, and brown=4. The logistic model was coded as follows: blue=1 versus non-blue=0 (brown and hazel). The linear model used for categorical eye colours followed an additive framework, wherein the estimated effect size corresponds linearly to the count of effect alleles.

The GWAS model included covariates such as age, sex, and the first ten principal components as fixed effects and a Genetic Relationship Matrix (GRM) as a random effect, alongside imputed genotype data.

### Meta-analysis

A meta-analysis was carried out using the beta coefficients and standard errors from the linear GWAS results corresponding to the two distinct CanPath arrays, along with data from each logistic GWAS using the software Metasoft (version 2.0.1) (22). Metasoft assesses input GWAS using two types of models, 1) fixed effects (FE), most appropriate when effect sizes are assumed to be consistent across studies, and 2) an enhanced random effects model (RE2), which is more suitable when there is variability or heterogeneity between datasets (23).

Metasoft computes two key indicators of heterogeneity: Cochran’s Q and I². The Q statistic reflects the weighted squared differences between individual study effects and the combined effect estimate; a P-value below 0.10 typically indicates significant heterogeneity. The I² metric (24,25) estimates the proportion of variation across studies that exceeds what would be expected by chance, calculated as I² = 100% × (Q – df)/Q.

When I² exceeds 50%, this may suggest that a random effects model is more appropriate than a fixed effects model (26,27). In the absence of heterogeneity, both models are expected to produce similar outcomes. Metasoft also generates Bayesian posterior probabilities indicating the likelihood that an association exists in each study included in the meta-analysis (23).

### Conditional and Joint Analysis (COJO) Using GWAS Summary Statistics and Fine-Mapping

The combined summary statistics from the linear GWAS meta-analysis were subsequently processed using the COJO method included in the GCTA suite (28). COJO is used to determine whether genome-wide significant signals (P-value<5E-8) identified in a GWAS (or meta-analysis) are attributable to a single variant or represent multiple independent signals. The analysis was based on fixed-effects (FE) model P-values from the meta-analysis and incorporated weighted average effect allele frequencies derived from both contributing datasets. A linkage disequilibrium (LD) reference was generated from the GSA array as it had the larger sample size of the two arrays for AA and AG individuals.

Statistical fine-mapping was performed using LDSTORE2 (v2.0) and FINEMAP (v1.4) (29), focusing on regions highlighted by the COJO analysis. This approach helps pinpoint variants most likely to causally influence eye colour. FINEMAP uses a Bayesian model to integrate GWAS or meta-analysis summary statistics with SNP LD data, producing posterior probabilities for putative causal variants.

In the first step, LDSTORE2 was used to generate LD correlation matrices from the GSA 24v1MDP array data within 1 Mb windows (±500 kb from lead SNPs). Next, these LD matrices were input into FINEMAP along with the defined genomic windows. The analysis assumed up to 10 causal variants per region, generating 95% credible sets – groups of SNPs with the highest posterior probability of being causal. Each set is summarized by its posterior inclusion probabilities and a k-value indicating the most probable number of causal variants.

### SNP Annotation

To annotate the genome-wide significant variants (P<5E-08) identified through meta-analysis, we employed SNPnexus, a web-based annotation tool (https://www.snp-nexus.org/v4/) (30–34). Gene and variant classification were based on data from the UCSC Genome Browser and Ensembl databases using the hg19 human genome build. If, through SNPNexus, we were able to determine if a variant was listed in ClinVar (35), then the clinical annotation was also documented. The potential functional impact of non-synonymous coding variants was evaluated using SIFT (36) and PolyPhen (37) prediction scores, which provide qualitative outputs such as “probably damaging”, “possibly damaging/deleterious-low confidence”, or “benign/tolerated”. For non-coding variants, we applied the CADD scoring system (38), which ranks variant deleteriousness across the genome. A CADD score of 20 or higher suggests the variant is among the top 1% of the most deleterious substitutions (34). Furthermore, we investigated protein-level expression of implicated genes using STRING (39) and searched for eQTLs (expression Quantitative Trait Loci) in GTEx (https://www.gtexportal.org/home/) (40).

### Replication in Independent Sample of Individuals of European Ancestry with Quantitative Measures of Iris Colour

The SNPs identified in the CanPath GWAS were replicated in a sample of individuals of European ancestry living in Canada, for whom detailed eye colour quantitative measures based on high-resolution iris pictures are available. The sample and the methods used to measure iris pigmentation have been described in a previous paper (41). The sample includes 549 individuals with ancestry from different regions in Europe. High-resolution iris pictures were taken using a Miles Professional Iris Camera and quantitative measures of iris colour using the CIE 1976 L*a*b* (CIELAB) colour space were obtained using a web-based application (41). The L* coordinate represents the lightness dimension and ranges from 0 to 100, with 0 being black and 100 being white. The a* and b* coordinates represent variation in colour, with negative values of a* indicating green and positive values of a* indicating red, and negative values of b* indicating blue and positive values of b* indicating yellow. Individuals with blue eye colour are characterized by negative a* and b* values, and higher L* values than those observed for individuals with non-blue eyes. However, there is considerably variation in L*, a* and b* dimensions within individuals reporting “blue” or “brown” eye colour (41). In addition to L*, a* and b* values, the amount of central heterochromia (i.e., differences in iris colour between the inner and outer portions of the iris) was measured by a colour metric (ΔE) that represents differences in the average colour of the ciliary zone (outer iris region) and the average colour of the pupillary zone Inner iris region). Two different cameras were used during the study, and camera was used as a covariate in the statistical analysis. Participants in the study also self-reported eye colour in 5 categories: 0=Light blue, Light gray or light green; 1=Blue, gray or green, 2=Hazel or light brown, 3=Dark brown, 4=Brownish black.

Genotyping was carried out with Illumina’s Infinium Multi-Ethnic Global Array (MEGA) at the Clinical Genomics Centre (Mount Sinai Hospital, Toronto, Ontario, Canada) using standard protocols. QC steps were implemented to remove samples and markers, according to the following criteria, Sample QC: 1) Removal of samples with missing call rates <0.9, 2) Removal of samples that were outliers in Principal Component Analysis (PCA) plots, 3) Removal of samples with sex discrepancies, 4) Removal of samples that were outliers for heterozygosity, and 5) Removal of related individuals (pi-hat> 0.2). Marker QC: 1) Removal of markers with genotype call rate <0.95, 2/removal of markers with Hardy-Weinberg Es <10−6, 3) Removal of Insertion/Deletion (Indel) markers, 4) Removal of markers with allele frequencies <0.01, 5) Removal of markers not present in the 1000 Genomes reference panel, or that do not match on chromosome, position and alleles, 6) Removal of A/T or G/C SNPs with MAF >40% in the 1000 Genomes European reference samples, and 7) Removal of SNPs with allele frequency differences >20% between the study sample and the 1000 Genomes European reference sample. After performing the QC steps described above, the samples were phased using the program SHAPEIT2 and imputed at the Sanger Imputation Service, using the Positional Burrows-Wheeler Transform (PBWT) algorithm (42), and the samples of the 1000 Genomes as reference haplotypes. All the markers included in this analysis have imputation INFO scores higher than 0.9, indicating good imputation confidence.

Following the same approach used for the CanPath sample, individuals were stratified in two categories based on *HERC2* rs12913832 genotype (GG and AA+AG). Association of the selected markers with the quantitative traits were tested using linear regression with sex, camera, and 4 principal components as covariates. Age was not included as a covariate because there was a narrow range in participant’s age (18-35 years) and age did not have a significant effect on the traits. Given that some quantitative traits (a*, b* and ΔE) showed non-normal distributions, an inverse normal transformation of the residuals from a regression of a*, b* and ΔE on sex, camera and 4 principal components was carried out, and these normalized values were used as dependent variables in the linear regression. A linear model was also used to evaluate association of the markers with the five self-reported eye colour categories (0=Light blue, Light gray or light green; 1=Blue, gray or green, 2=Hazel or light brown, 3=Dark brown, 4=Brownish black).

## Results

### Preliminary Statistics and Quality Control (QC)

After applying all QC filters, genotypes and self-reported eye colour phenotypes were retained from a total of 2,724 AA+AG individuals (1,412 from the GSA array and 1,312 from the UKBB array). Eye colour categories included blue, green, hazel and brown. Of the 2,724 rs12913832:AA+AG participants, 512 did not self-report brown or hazel eyes, representing 41.16%. As was reported in 2023, of the 2,757 rs12913832:GG participants for which self-reported eye colour data were available, 904 did not report blue eyes. **Fig 4** illustrates eye colour frequency stratified by genotype at the rs12913832 locus. In total, 5,481 CanPath participants are represented when considering all rs12913832 genotypes combined.

**Fig 4.**
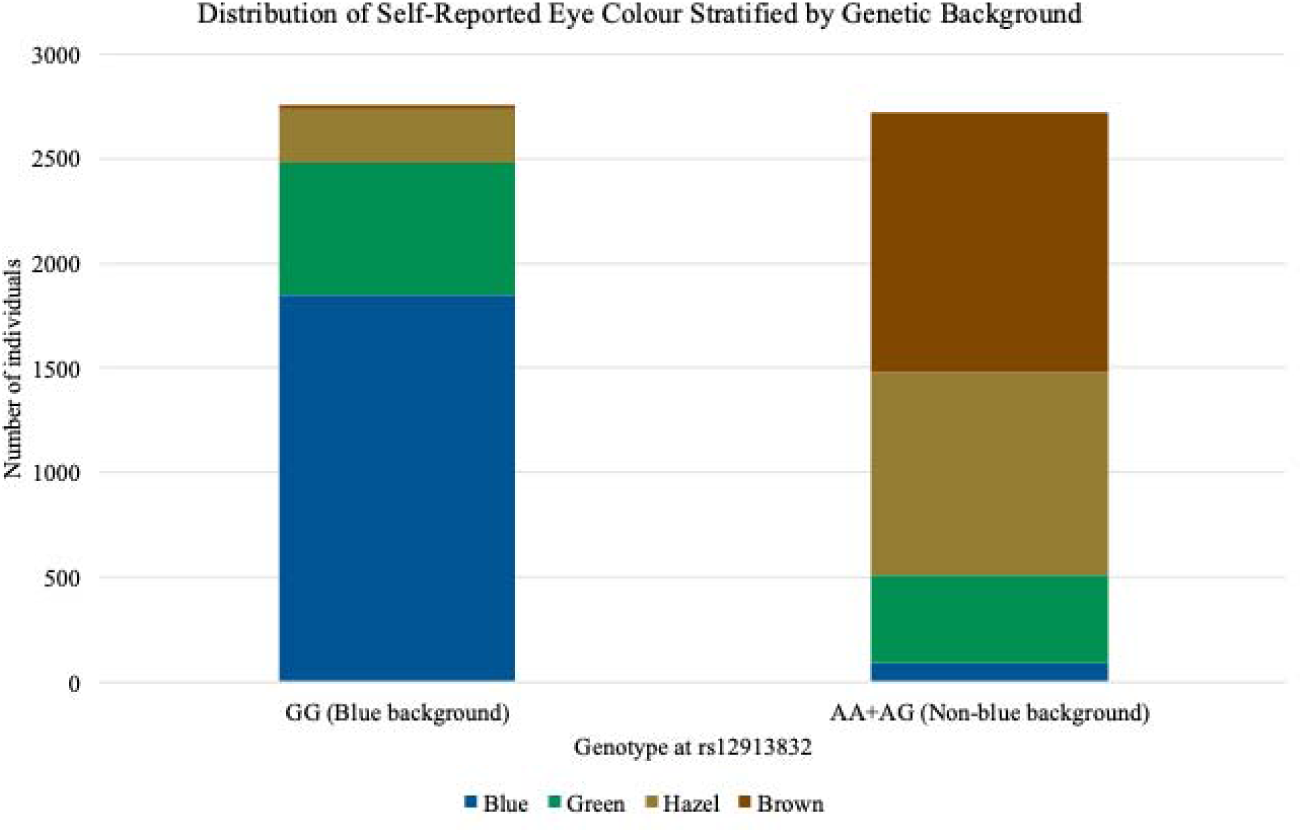
Eye colour distribution among 2,757 rs12913832:GG and 2,724 rs12913832:AA+AG Canadian Partnership for Tomorrow’s Health (CanPath) participants genotyped with the UKBB array and GSA array. Self-reported eye colour categories retained for investigation were described as blue, green, hazel, and brown. Individuals with missing phenotype information or who self-reported their eye colour as amber were excluded.

After QC and imputation 8,822,293 variants were retained from AA+AG individuals genotyped with the GSA array for further analysis and 7,104,980 variants were retained from AA+AG individuals genotyped with the UKBB array.

### Genome-Wide Association Study (GWAS) and Meta-Analysis

As was performed on the GG individuals in the 2023 study, two types of GWAS were performed on AA+AG individuals from each array using SAIGE (v1.0.5) (21). A linear mixed model was applied where eye colour was coded as blue=1, green=2, hazel=3, and brown=4 in accordance with an additive genetic model, as well as a logistic model comparing blue versus non-blue (hazel and brown) eye colour. The first ten genetic principal components generated during the QC stage were included as covariates, along with age and sex.

The summary statistics (P-values, betas, and SE) from the linear and logistic models conducted in each CanPath array GWAS were then integrated in a meta-analysis using Metasoft (v2.0.1) (22). The resulting Manhattan plot from the linear model is illustrated in **Fig 5** (Supplementary **Figs S1** and **S2** present the QQ plots for each meta-analysis and the Manhattan plot from the logistic model). Significant peaks are observed in chromosomes 6 (*IRF4*), 9 (*TYRP1*) and 15 (*OCA2*).

**Fig 5.**
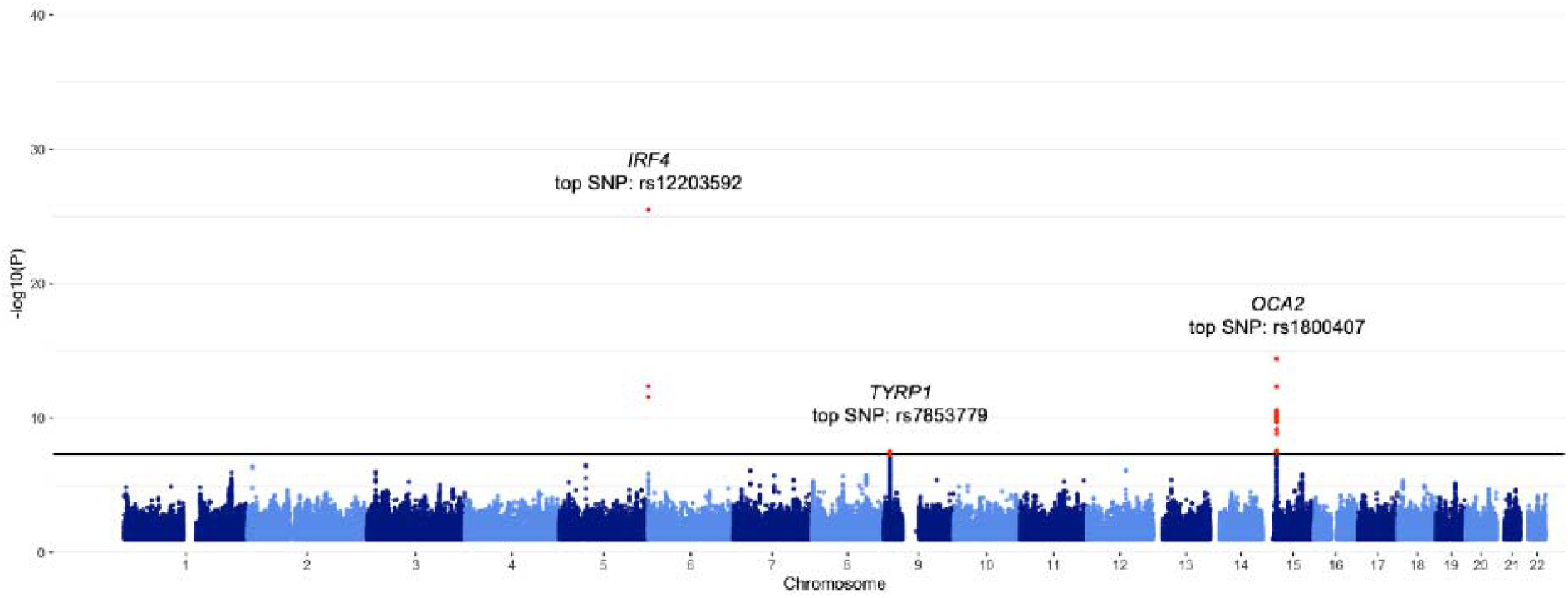
Manhattan plot of eye colour meta-analysis based on linear model GWAS in rs12913832:AA+AG individuals (*N*=2,724) from the Canadian Partnership for Tomorrow’s Health (CanPath) project genotyped with the Axiom UKBB and GSA arrays. GWAS were performed with SAIGE (21). Categorical eye colours coded as blue=1, green=2, hazel=3, and brown=4. The meta-analysis was conducted with Metasoft (22). The black horizontal line indicates the genome-wide threshold (P<5E-8). Significant peaks are observed in chromosomes 6 (*IRF4*), 9 (*TYRP1*) and 15 (*OCA2*).

### Comparing GWAS Lead SNPs from GG and AA+AG Individuals

**Table 1** illustrates the top SNPs from the meta-analysis according to the fixed effects (FE) linear model in each significant genomic region. Several SNPs had evidence of heterogeneity among the two studies (indicated by I^2^ and Cochran’s Q values in the meta-analysis), however, they all retained genome-wide significant P-values in the random effects (RE2) model too, which considers heterogeneity among studies. Given that heterogeneity between studies did not impact SNP significance, the fixed effects (FE) model was prioritized for downstream analyses. Please note that that in the 2023 publication (12) the effect alleles in the tables were incorrect, and a correction was submitted.

**Table 1.** Top SNP from each significant genomic region in meta-analysis of GG stratified individuals and AA+AG stratified individuals using a linear model (*N_GG_*=2,747, *N_AA+AG_*=2,724).

| Marker ID | Chr | A1 | A2 | A1<br>Frequency | Gene | P-value<br>(FE) | Beta | SE |
| --- | --- | --- | --- | --- | --- | --- | --- | --- |
| **GG Stratified Individuals** |  |  |  |  |  |  |  |  |
| **rs35389** | 5 | **G** | A | 0.046 | *SLC45A2* | 1.18E-12 | 0.352 | 0.049 |
| **rs2733836** | 9 | **T** | G | 0.438 | *TYRP1* | 1.37E-13 | 0.133 | 0.018 |
| **rs1126809** | 11 | A | **G** | 0.272 | *TYR* | 4.58E-13 | -0.179 | 0.019 |
| **rs12896399** | 14 | T | **G** | 0.401 | *SLC24A4* | 2.05E-35 | -0.221 | 0.018 |
| **rs62075722** | 17 | A | **G** | 0.364 | *TSPAN10* | 6.36E-10 | -0.116 | 0.019 |
| **AA+AG Stratified Individuals** |  |  |  |  |  |  |  |  |
| **rs12203592** | 6 | **T** | C | 0.164 | *IRF4* | 3.13E-26 | -0.318 | 0.030 |
| **rs7853779** | 9 | A | **G** | 0.389 | *TYRP1* | 3.12E-8 | 0.129 | 0.023 |
| **rs1800407** | 15 | **T** | C | 0.128 | *OCA2* | 4.10E-15 | -0.292 | 0.037 |

Based on the categorical coding of the model (1=blue, 2=green, 3=hazel, 4=brown), a positive beta can be interpreted as an indication of eye colour “darkening” and a negative beta can be interpreted as an indication of eye colour “lightening” for the effect allele (A1). We highlight alleles increasing pigmentation in the rs12913832 GG (e.g. blue-eye) background with red font and alleles decreasing pigmentation in the rs12913832 AA+AG (e.g. non blue-eye) background in blue font.
**Marker ID**, reference SNP cluster ID;
**Chr**, chromosome;
**A1**, effect allele: the allele whose effects in relation to eye colour are being studied;
**A2**, the alternative allele;
**A1 frequency**, fraction of chromosomes in population that carry the effect allele;
**Gene**, the gene the SNP is located in, or the gene the SNP is nearest to;
**Beta**, per unit increase (if positive) or decrease (if negative) in the outcome;
**P-value (FE)**, parameter of statistical significance used to determine the certainty of an association, based off the fixed effects meta-analysis model;

### Conditional and Joint Analysis (COJO)

Results from the COJO analysis using GCTA (28) for the GG individuals were previously described in 2023 (12), but they are summarized in **Table 2** to easily compare the different signals coming from each genetic background. The COJO analysis of the linear model conducted on AA+AG individuals identified one significant genome-wide SNP driving the signals on chromosomes 6 and 9, while two genome-wide significant SNPs were observed driving the signal on chromosome 15. Among these, rs12203592 in *IRF4* is a well-established causal pigmentation variant, previously associated with both skin and hair pigmentation, as well as melanoma susceptibility (6,43,44). The SNP rs7853779 on chromosome 9, located near *TYRP1*, had prior associations with pigmentation traits and melanoma susceptibility (1,45). Two SNPs on chromosome 15, rs1800407 and rs4778138, both situated in or near the *OCA2* gene, have been linked to variation in eye colour in earlier research (46–48).

**Table 2.**
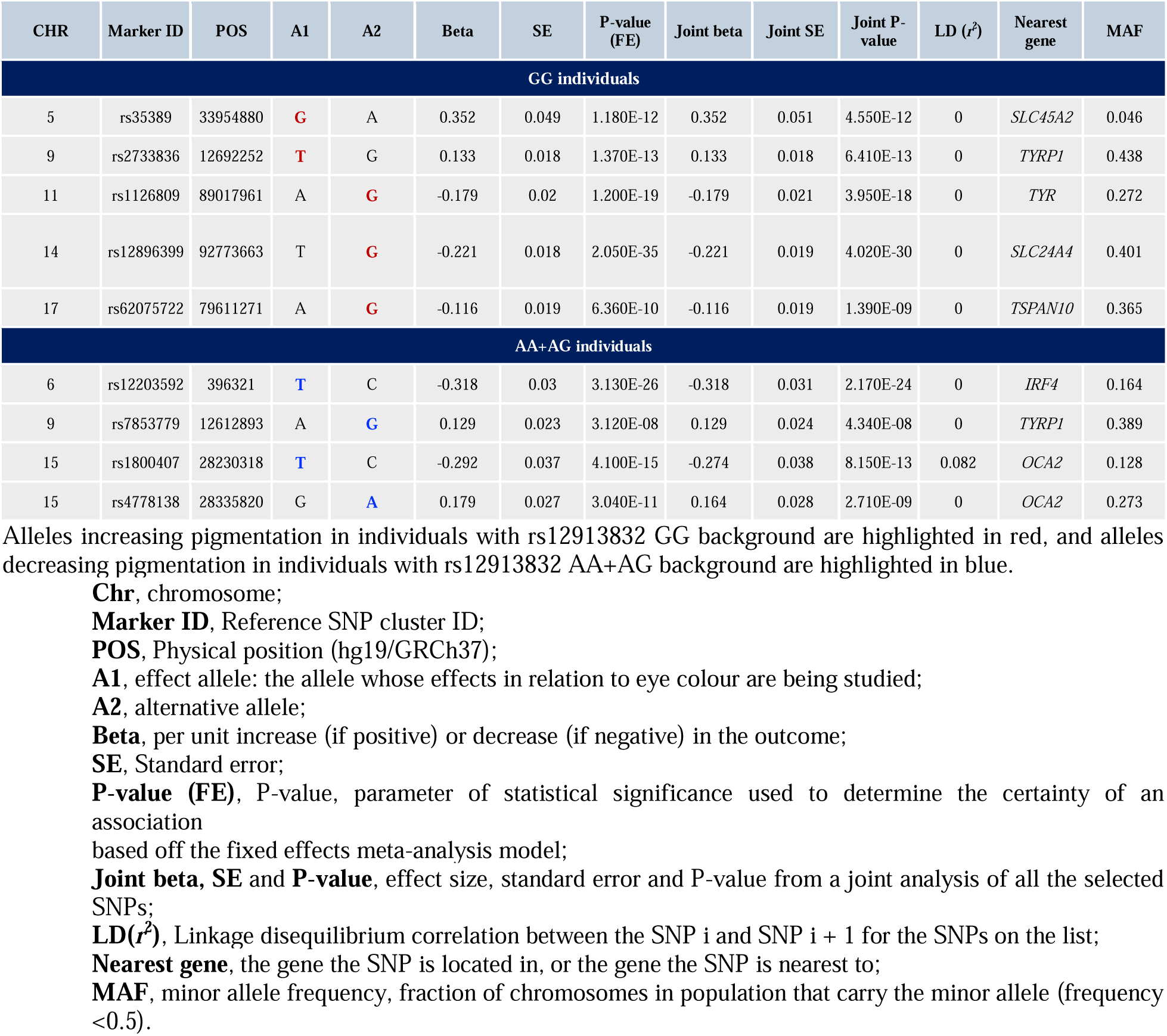
Independent signals identified by COJO (28) for genome-wide significant peaks illustrated in the meta-analysis.

### Fine-mapping

Fine-mapping results are represented in **Table 3**. Only SNPs with log_10_ of Bayes Factor (log_10_BF) greater than 2 have been included in the table. For some regions (*TYRP1*, *TYR*, *SLC24A4* and *TSPAN10* for the GG stratified individuals), the *k* value (most likely number of causal SNPs) is greater than one, however, only SNPs within one credible set surpassed a log_10_BF>2. Thus, although FINEMAP suggests the presence of multiple signals in some instances, the markers in the additional credible sets did not always have sufficiently high log_10_BF, so we focused on the markers that surpassed the thresholds for “considerable evidence” as recommended by the FINEMAP documentation (29).

**Table 3.**
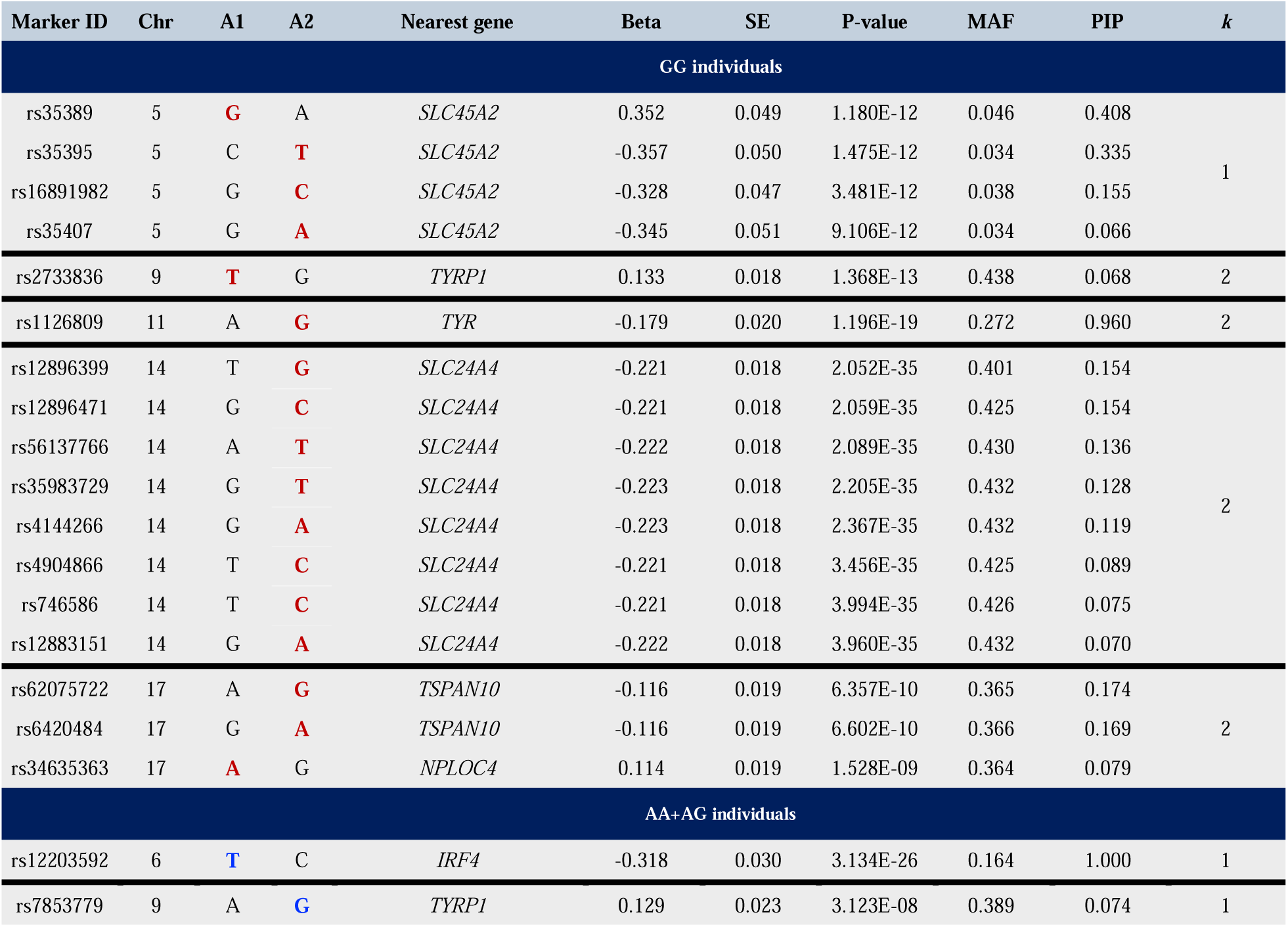
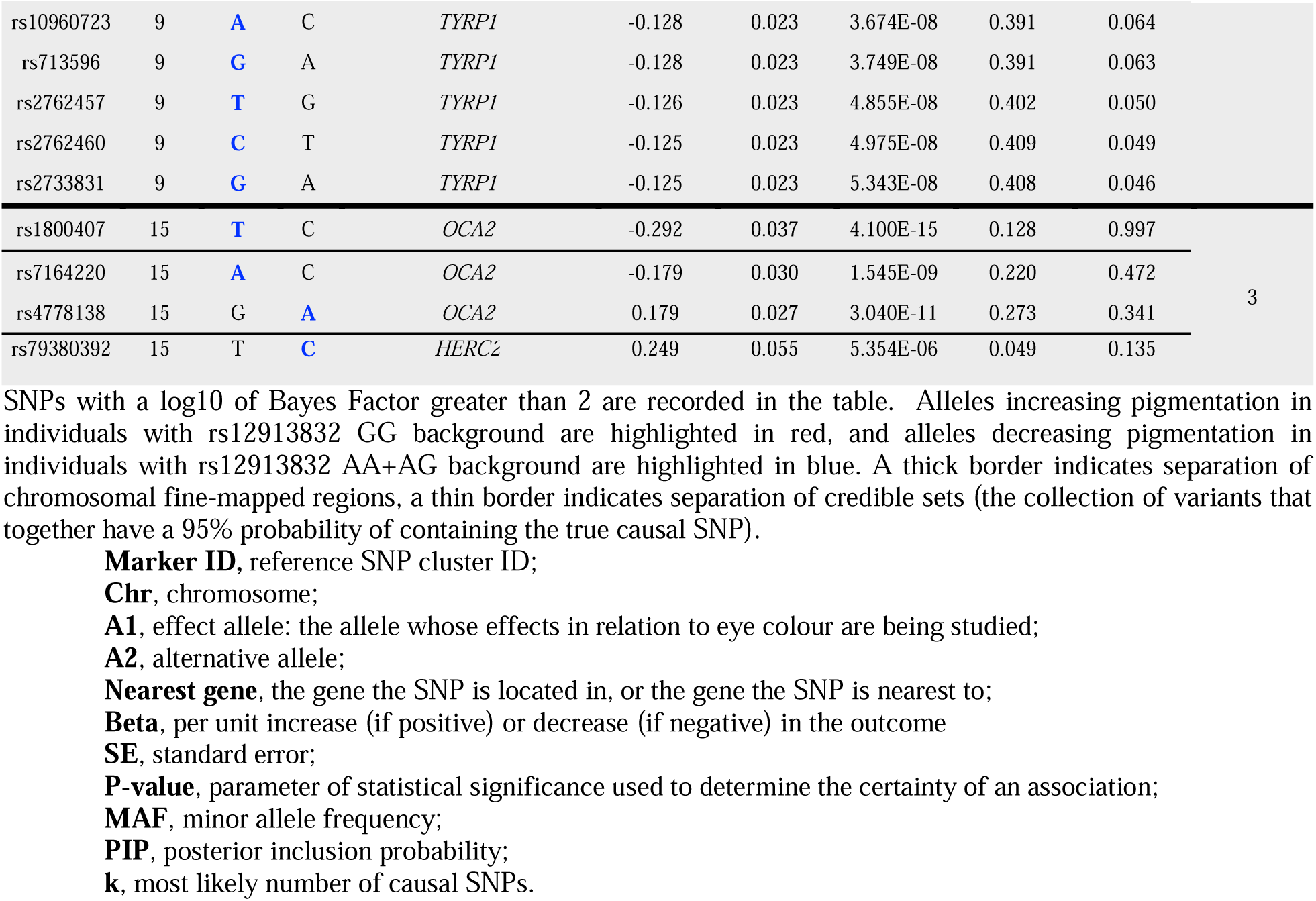
Fine-mapping results. Fine-mapping was performed using 1Mb windows (500 kb on each side of significant SNP).

For the GG stratified individuals, the markers with the top PIP in the FINEMAP analysis coincide with the lead SNPs identified in the COJO analysis: *SLC45A2* rs35389, *TYRP1* rs2733836, *TYR* rs1126809, *SLC24A4* rs12896399 (there is another marker with the same PIP in this region, rs12896471), and *TSPAN10* rs62075722. The non-synonymous variant rs1126809 is highlighted as the causal variant in the *TYR* region, with PIP=0.960. All the other variants have PIP values lower than 0.5.

For the AA+AG stratified individuals, the FINEMAP analysis identifies a single credible set in the *IRF4* and *TYRP1* regions. The marker rs12203592 is highlighted as the putative causal SNP within the *IRF4* region, with PIP=1.00. In contrast, there are several markers in high LD in the credible set of the *TYRP1* region, with PIP values<0.10. The top SNPs identified by FINEMAP coincide with the lead signals in the COJO analysis. Finally, the FINEMAP analysis identifies 3 independent credible sets in the *OCA2/HERC2* region. In the first credible set, FINEMAP identifies rs1800407 as the causal variant, with PIP=0.997. In the second credible set, FINEMAP highlights two SNPs: rs7164220 (PIP=0.472) and rs4778138 (PIP=0.341). The latter was the lead SNP identified in the COJO analysis. The variant with the top PIP in the third credible set is rs79380392 (PIP=0.135), which is located within the *HERC2* gene. In contrast to FINEMAP, which points to three independent causal sets, COJO only identified two independent regions in the *OCA2/HERC2* region. **Fig S3** illustrates LD matrices for SNPs in the *TYRP1* region and *HERC2/OCA2* region. **Fig S4** illustrates examples of regional plots for signals represented in **Table 3**.

### SNP Annotations

Beyond the rs12913832 genotype, acting as a genetic background in this study, three other SNPs within *OCA2* and one within *HERC2* were nominated as candidate causal loci in AA+AG stratified individuals. One of them is a non-synonymous SNP (rs1800407) and three are intronic variants (rs4778138, rs7164220, rs79380392). These results are similar to the conditional analysis with GCTA-COJO, where the program picked up multiple independent signals coming from this region. We annotated the putative regulatory function of the fine-mapped SNPs with a log_10_ of Bayes Factor greater than 2 using diverse databases (e.g. ENCODE, Roadmap Epigenomics Project) encompassed within SNPnexus (30–34). A summary of the output can be found in Supplementary **Table S1**. All SNPs in the table were identified as either overlapping with a protein-coding gene or were located near a protein coding gene by SNPNexus (30–34). Annotation details for the lead SNPs identified by COJO are described in **Table 4** below.

**Table 4.**
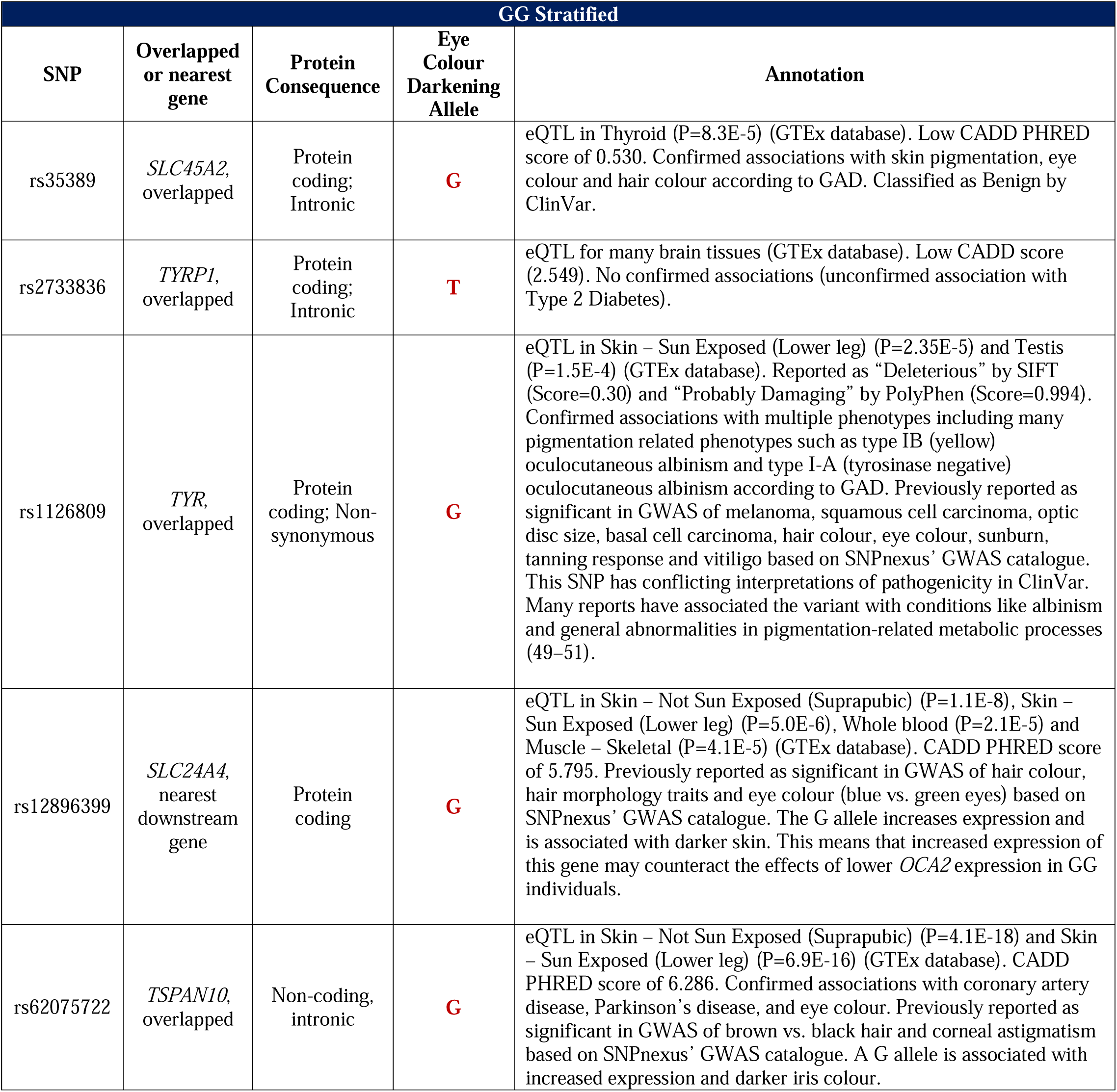
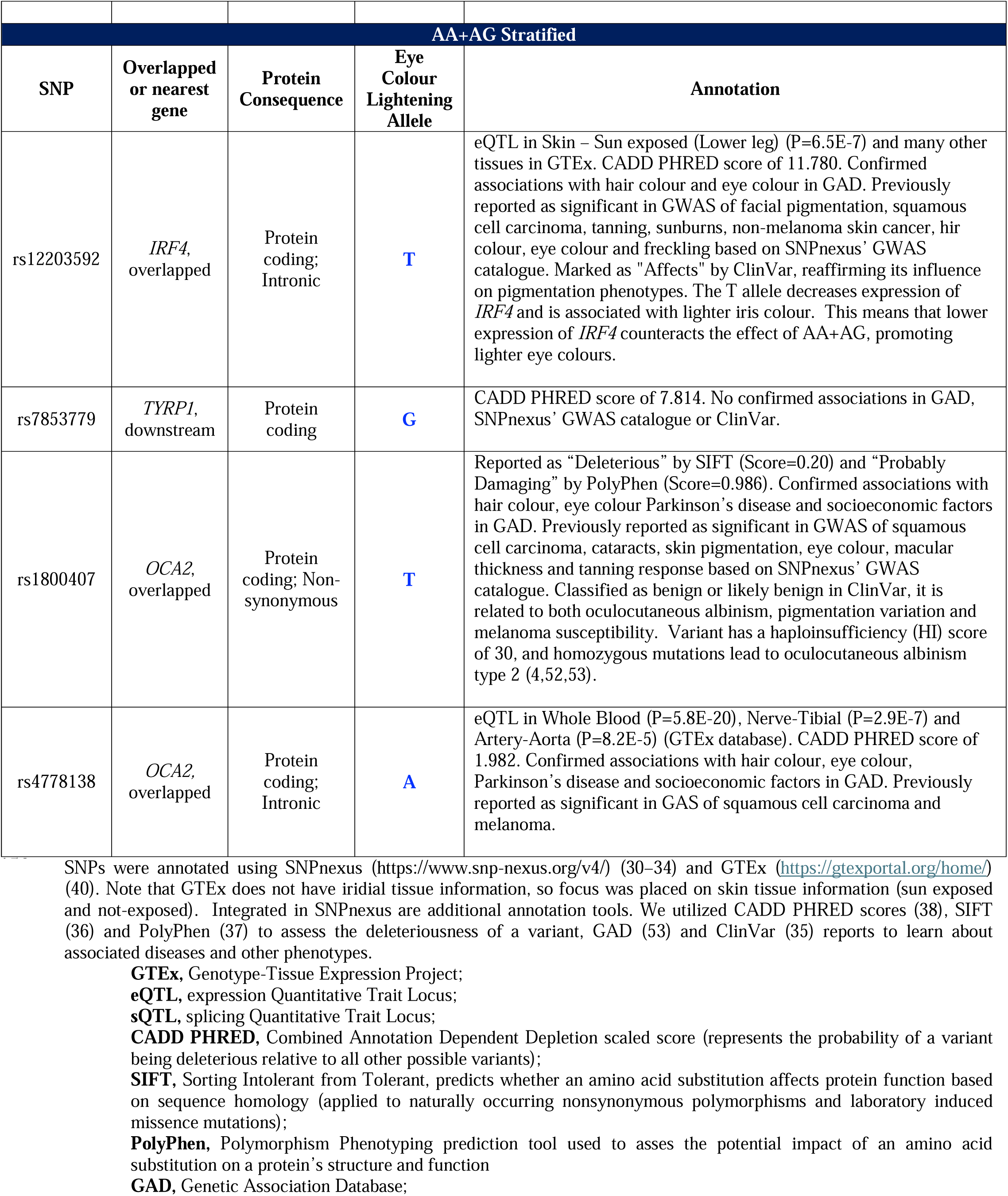
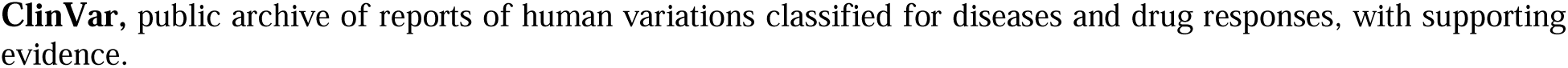
Annotation summary of lead SNPs.

Supplementary **Fig S5** and **Table S2** illustrate the STRING network and summarize the functional enrichments within the networks. There are slight differences in the biological processes enriched for genes which were associated with GG individuals versus AA+AG individuals. In GG individuals, genes can be seen enriched for melanin biosynthetic processes and developmental pigmentation, while in AA+AG individuals, genes were enriched in melanosome membrane. Interestingly, under the “Human Phenotype (Monarch)” STRING domain, the genes associated with the AA+AG, or brown eye background, the phenotype with the strongest signal (FDR=1.96E-5) was blue irides, or a markedly blue colouration of the iris.

### Replication in European ancestry cohort with quantitative iris pigmentation measures

Replication results for the six different analyses are reported in Supplementary **Table S3** (Supplementary **Fig S6** illustrates the distributions of rs12913832 genotype across self-reported eye colour category for this replication cohort). Of the five independent signals from GG individuals, three markers survived Bonferroni correction in at least one quantitative measure (*TYR* rs1126809 in the L and b* dimensions, *SLC24A4* rs12896399 in the b* dimension, and *TSPAN10* rs62075722 in the b* dimension). Of the four independent signals from AA+AG individuals, three markers passed Bonferroni correction in multiple dimensions (*IRF4* rs12203592 in the L, a*, b* and ΔE dimensions, *OCA2* rs1800407 in the L and a* dimensions, and *OCA2* rs4778138 in the L and a* dimensions). Two markers demonstrated nominal significance in other quantitative measurements (*SLC24A4* rs12896399 for the GG stratified sample and *TYRP1* rs7853779 for the AA+AG stratified sample,). The effects appear strongest in the b* dimension (Blue [-] to yellow [+]) for GG stratified individuals and in the L dimension (Lightness dimension: 0 (black) to 100 (white)) for AA+AG stratified individuals. We also conducted a linear regression using the self-reported eye colour categories in the independent replication dataset, and the results were concordant with the quantitative measures. In GG stratified individuals, *TYR* rs1126809 was nominally significant, and in AA+AG stratified individuals, all markers, with the exception of *TYRP1* rs7853779, survived Bonferroni correction. Importantly, rs12913832 was also assessed in the replication dataset in AA+AG stratified individuals. This marker illustrated significance in the L, a* and ΔE dimensions, as well as the linear model – indicative that eye colour is dosage dependent with respect to genotype at rs12913832. In particular, rs12913832 is very strongly associated with ΔE (p=2.97E-07), suggesting that having one copy of the G allele increases the likelihood of central heterochromia (e.g. differences in iris colour between the inner and outer rings of the pupil).

## Discussion

This study expands upon previous work investigating genetic modifiers of iris pigmentation by analyzing individuals whose self-reported eye colour does not align with the expected phenotype based on their *HERC2* rs12913832 genotype. Specifically, we focused on those with a homozygous derived genotype (rs12913832:GG), which typically predicts blue eyes, and heterozygous or homozygous ancestral genotypes (rs12913832:AG or AA), which are more often associated with brown eyes. Despite the heavy reliance on this single SNP in forensic and anthropological models, exceptions in phenotype suggest additional modifiers. Our genome-wide approach successfully identified several SNPs that modify eye colour in these “non-concordant” individuals and replicated key findings using high-resolution quantitative iris colour data in an independent European sample.

Through our GWAS meta-analyses and fine-mapping analyses we identified loci in known pigmentation-associated regions including *SLC45A2*, *TYRP1*, *TYR*, *SLC24A4*, and *TSPAN10* in the GG group (where blue eyes is expected) and *IRF4*, *TYRP1*, and *OCA2* in the AA+AG group (where brown eyes is expected).

### Genotype-stratified Associations with Eye Colour in Recent Studies

In 2020, Meyer and colleagues (10) conducted a candidate gene study investigating the association between brown eye colour in rs12913832:GG individuals and SNPs in previously known pigmentation genes (*TYR*, *TYRP1* and *SLC24A4*). The group identified four variants to be the most promising candidates for the explanation of the genotype-phenotype discordances observed in GG individuals: *TYRP1* rs35866166:C, *TYRP1* rs62538956:C, *SLC24A4* rs1289469:C, and *TYR* rs1126809:G. Of these four markers, three were replicated in our fine-map analyses: *TYRP1* rs35866166:C, *TYRP1* rs62538956:C and *TYR* rs1126809:G. The *TYR* rs1126809 SNP is represented in **Table 3** and has a high PIP of 0.96, strongly suggesting it is the causal variant, with the G allele strongly associated with darker iris pigmentation in rs12913832:GG individuals. The *TYRP1* rs35866166 and *TYRP1* rs62538956 markers are present in our 95% credible set, however, they each had very low PIP values (0.001 and 0.003 respectively) and our fine-mapping analysis prioritize a different variant within this region (rs2733836). That being said, we did find that the effect direction was the same in our study, with the rs35866166:C and rs62538956:C alleles corresponding to positive beta values (in the direction of eye colour darkening with our linear model). The authors did note that none of the variants identified in the candidate gene study showed statistically significant associations with eye colour after traditional multiple testing correction, but in our GWAS we found rs1126809 and rs62538956 to surpass genome-wide significance (P=1.200E-19 and P=5.846E-9 respectively), and rs35866166 to be nominally significant (P=2.950E-6). Three years later, in 2023, Salvo and colleagues (54) performed an additional candidate gene study investigating the association between variants in the *HERC2/OCA2* region and blue eye colour in rs12913832:AA+AG individuals. They identified five novel variants to be the most promising candidates for explanation of blue eye colour in individuals with a brown-eye background: rs74409036:A (*OCA2*), rs78544415:T (*OCA2*), rs72714116:T (*OCA2*), rs191109490:C (*HERC2*) and rs551217952:C (*HERC2*). None of these SNPs are genome-wide significant in our analysis, and our analysis points to different markers within this region. We describe in more detail the main regions and polymorphisms identified in our stratified analysis below.

### Eye Colour Darkening in GG Stratified Individuals

#### SLC24A4 and SLC45A2

*SLC24A4* encodes a sodium/calcium/potassium exchange protein (GeneCards, 2025). Variants in this gene have been previously implicated in skin, hair and eye pigmentation, although its precise function in melanogenesis remains unclear (55,56). Our lead SNP in this region is rs12896399, which is situated approximately 15,000 base pairs upstream of the *SLC24A4* transcription start site and has been included in established eye colour prediction tools, including IrisPlex (57–59) and Snipper 3.5 (60). The T allele of rs12896399 is associated with blue eye colour (59), a relationship also observed in this study. In our preliminary 2023 investigation of rs12913832:GG individuals (12), we observed strong linkage disequilibrium among SNPs within the *SLC24A4* credible set, with pairwise comparisons yielding *r*² values greater than 0.9. eQTL data in GTEx indicates that a G allele at rs12896399 is associated with increased *SLC24A4* expression, suggesting that higher expression of this transporter is able to compensate for the reduced expression of *OCA2* in rs12913832:GG individuals. Our results point to rs12896399:G, or closely linked variants, as strong candidates for darker eye colours in GG individuals.

*SLC45A2* encodes another transmembrane transporter involved in melanin biosynthesis. This gene is expressed in melanoma cell lines and is implicated in oculocutaneous albinism type 4. Its polymorphisms are most commonly associated with variation in skin and hair pigmentation (61). The *SLC45A2* variant rs16891982 is included in both the IrisPlex (57–59) and The Snipper 3.5 (60) models, and appeared in our COJO and fine-mapping analyses of rs12913832:GG individuals. However, rs16891982 was not the top candidate within the credible set (PIP=0.155). Instead, rs35389 emerged as the most likely causal variant, with a PIP of 0.408. These two SNPs are in linkage disequilibrium, as confirmed by LDpair (D’=1.0, R²=0.6964, χ²=700.577, P<0.001 with EUR reference) (Machiela & Chanock, 2015). While rs35389 has not been previously associated with eye colour, it has been evaluated as an ancestry informative marker (62) and has shown significant associations with methylation patterns in the context of population stratification (63).

#### TYRP1

Notably, *TYRP1* emerged as a consistent locus across both genetic backgrounds. *TYRP1* encodes the tyrosinase-related protein 1, which is a member of the tyrosinase family and is involved in the synthesis of eumelanin (64). Specifically, it is responsible for the stabilization of tyrosinase, which is an important enzyme in the initiation of melanin synthesis (65) and contributes to melanosomal structure and maturation (66). *TYRP1* is primarily expressed in melanocytes and the retinal pigment epithelium (RPE) (67). Only one marker, rs2733836, illustrated association in the GG background. There are no *TYRP1* variants currently included in eye colour prediction tools specifically, although rs683 appears for the prediction of both hair and eye colour (57–59). In 2009, Liu et al. (55) compared predictive power of a six-SNP model (*HERC2* rs12913832, *OCA2* rs1800407, *SLC24A4* rs12896399, *SLC45A2* rs16891982, *TYR* rs1393350, and *IRF4* rs122203592) with a 15 SNP model where two *TYRP1* variants were included (rs1408799 and rs683) and found that the inclusion of the additional SNPs provided very minimal additive effects and did not significantly improve predictive accuracy. rs2733836 (associated in this study), rs1408799 and rs683 are in linkage disequilibrium (D’>0.7) (Supplementary **Fig S3**), yet rs2733836 has not previously been associated with eye colour. Not much is known of the variant’s consequence apart from the fact that it is located 2 kilobases upstream of *TYRP1*; the variant itself is not a protein coding variant and is therefore not expressed, but the gene is, and can be found most highly expressed in skin (30–34).

#### TYR

The primary *TYR* variant identified in our study was rs1126809, a coding SNP in exon 4 of *TYR,* and most likely causal variant in this region (PIP=0.960). The alternative allele (A) induces a missense mutation that renders the tyrosinase enzyme thermosensitive and less efficient, reducing melanin synthesis (68,69). Given that *TYR* is the rate-limiting enzyme in melanogenesis (70,71), individuals carrying the A allele are expected to exhibit lighter pigmentation. Our results indicate that the presence of the rs1126809 G allele can oppose/counteract the effects of reduced *OCA2* expression driven by the rs12913832:GG genotype

There is currently only one *TYR* variant included in eye colour phenotyping models: rs1393350 (57–60,72,72–76). rs1393350 and rs1126809 are in near-perfect linkage disequilibrium (D’=1.0, R²=0.9532, χ^2^=958.8784, P<0.001, with EUR reference), as reported by LDpair (77). Our findings echo that of Meyer et al., 2020 (10) and raise the possibility that rs1126809 may be a more biologically informative marker than rs1393350. Both variants reached genome-wide significance in our GWAS and were identified in fine-mapping analyses; however, rs1393350 demonstrated a notably low posterior inclusion probability (PIP=0.017) despite its strong P-value (P=5.81E-18), suggesting it may not be the causal variant but rather tags a signal in high LD. Importantly, while rs1393350 is known to associate with iris colour (with the A allele enriched in green eyes and the G allele in blue) (56), its moderating function remains poorly defined. In contrast, rs1126809 is a functional missense variant that reduces tyrosinase enzymatic activity through thermosensitivity, directly impacting melanin production.

#### TSPAN10

*TSPAN10*, or tetraspanin-10, has recently been associated with both quantitative and categorical measurements of eye colour (1,63). It is known to be expressed in the eye, including iris, ciliary body and RPE (78), however, its specific contribution to pigmentation variation has yet to be fully identified. There are no *TSPAN10* (or *NPLOC4*) variants currently included in any DNA phenotyping tools. In our analysis, rs62075722 illustrated the highest PIP (0.174) of the credible set and is an eQTL in GTEx (40), although it should be noted that a PIP of 0.174 is low and high LD among variants in the credible set makes it difficult to identify a singular putative causal variant. The expression data indicates that a G allele at this locus is associated with increased expression and a darker iris colour in GG stratified individuals. This is concordant with the other two variants identified by FINEMAP, rs6420284 and rs34635363, which are also significantly expressed in skin tissue in GTEx. Both the rs6420284:G allele and the rs34635363:G allele are associated with increased expression and darker iris pigmentation.

### Eye Colour Lightening in AA+AG Stratified Individuals

#### IRF4

The transcription factor *IRF4* has been shown to enhance *TYR* gene expression through interaction with MITF, a key regulator in melanocyte development (6). One specific *IRF4* variant, rs12203592, has been demonstrated to influence *IRF4* expression in melanocytes (79), and is included in the IrisPlex prediction panel (57,58). In the present study, we detected a significant association between eye colour and rs12203592 in individuals carrying the AA or AG genotypes at rs12913832. Prior research also suggests that interactions between *IRF4* and *OCA2* may modulate eye pigmentation. For instance, both Laino et al. (80) and Lona-Durazo et al. (11) reported that individuals with rs12913832 AA or AG genotypes – generally linked to darker eye colour – can still present with blue eyes if they carry one or two copies of the rs12203592:T allele, which has been associated with lighter pigmentation (80). The T allele at rs12203592 leads to decreased *IRF4* expression (40). Phenotypically, carriers of the T allele tend to exhibit lighter eye colour (e.g., blue or green), particularly among those with the rs12913832:AA or AG genotypes, where the effect of the T allele can partially override the darker eye phenotype typically associated with these genotypes.

#### TYRP1

The observation of *TYRP1* variants (rs7853779, rs10960723, rs713596, rs276257, rs2762460, rs2733831) in the AA+AG genetic background is both interesting, and more difficult to interpret as there are no other studies, candidate gene or otherwise, which have investigated the role of *TYRP1* SNPs in individuals with an rs12913832:AA+AG genetic background. Of the six *TYRP1* variants with a log10_BF_>2, only three have published associations in scientific literature. rs2762460 has been previously associated with breast cancer (81), and epidemiological studies have proposed that melanoma and breast cancer may share genetic factors (82,83), positioning *TYRP1* as a pleiotropic candidate gene. rs2762457 was used as a tag SNP in a candidate gene study of individuals with a blue eye genetic background, however, it was determined not to be associated with eye colour, but rather in strong LD with other SNPs associated with eye colour (10). Of note, our study indicates that variants within this gene, which is known to play an important role in melanin synthesis, may be responsible for darker eye colours in rs12913832:GG individuals and lighter eye colours in rs12913832:AA+AG individuals.

It is also interesting to note that the *TYRP1* variant identified in the GG genetic background is in high LD with the *TYRP1* variants identified in the AA+AG genetic background (all pairwise relationships have a D’>0.9 and R^2^>0.69, as confirmed by LDmatrix (with EUR reference) (77)). Variants in this region may make good candidates for SNP-SNP interaction tests in future studies, however, it would be important to consider that formal tests of epistatic interactions often suffer from statistical power and multiple hypothesis testing limitations.

#### HERC2/OCA2

*OCA2* encodes the P protein, a critical transmembrane component of the melanosome that plays a vital role in melanin biosynthesis and pigmentation (84). In addition to transporting melanosomal content, the P protein helps regulate melanosomal pH, which in turn influences both the amount and quality of melanin deposited in melanocytes (5,7). Dysfunction or loss of this protein, observed in both mice and humans, results in oculocutaneous albinism, underscoring its essential function in pigmentation pathways (7,85). Regulation of *OCA2* expression occurs via a long-range enhancer located within intron 86 of the neighbouring *HERC2* gene. Reduced expression of *OCA2*, mediated by variation in this enhancer, disrupts melanosome maturation and limits the effective processing and transport of melanin. This results in lighter pigmentation (such as that observed in blue eyes) due to compromised P protein availability (5).

In our study, multiple *HERC2/OCA2* variants were observed to be associated with eye colour variation in AA+AG individuals. By applying a Bayesian fine-map approach we found that the most likely number of causal variants for this genomic region was three, for which three candidates were located in *OCA2* (rs1800407, rs4778138/rs7164220) and one candidate in *HERC2* (rs79380392). rs1800407 is a missense variant currently included in forensic phenotyping tools (57–59). The rs1800407:T allele encodes the amino acid glutamate and having just one copy of the T allele is associated with an increased probability of having green and hazel eye colours in European populations (52,53). Given this, the presence of the T allele at rs1800407 in our study may help explain the elevated frequency of self-reported green and hazel eye colours by individuals with rs12913832:AA or AG genotypes (**Fig 4**). With a PIP of 0.997, and the only SNP with a log_10_BF>2 in this credible set, rs1800407 appears to be the first of three causal SNPs in this region. In the second credible set (rs7164220 and rs4778138), it is difficult to parse which variant is the causal variant. rs4778138 was identified as an independent signal in the COJO analysis, however, rs7164220 possesses the slightly higher PIP in the FINEMAP analysis (PIP=0.472). rs4778138 has been previously identified as being involved in a three-SNP haplotype system (rs4778138, rs4778241, and rs7495174) associated with blue eyes (86). In addition to rs4778138, we also observed rs7495174 from the blue eye haplotype in our study; although it was only nominally significant (PIP=004, P=1.87E-7), it may help explain the instances of blue eyes in the AA+AG cohort. Finally, rs79380392 appears to be the causal SNP in the third credible set as the only SNP with a log_10_BF>2. It is important to note that, although highlighted by FINEMAP, this SNP did not reach genome-wide significance in our study (Table 3), so this putative causal variant must be interpreted with caution.

### Replication

The replication analyses provided strong confirmation for the majority of associations identified in the discovery cohort. Among GG stratified individuals, three out of five independent signals (*TYR* rs1126809, *SLC24A4* rs12896399 and *TSPAN10* rs62075722) replicated with Bonferroni-corrected significance in at least one quantitative dimension. These associations were most commonly observed in the b* dimension (blue-yellow). Among AA+AG individuals, three of four loci (*IRF4* rs12203592, *OCA2* rs1800407, and *OCA2* rs4778138) survived Bonferroni correction across multiple quantitative measures. These associations were most commonly observed in the L* (lightness) dimension, consistent with the decrease in eumelanin concentration that accompanies a transition from brown eye colours to intermediate and blue eye colours. The results from the linear model using self-reported eye colour categories were also quite consistent with the CanPath sample results, with the replicated variants showing the same effect directions as those observed in linear CanPath analysis.

Importantly, rs12913832 was included in the AA+AG replication because there is variation in this group (1 versus 2 copies of the ancestral allele). The results indicate that having one derived allele (G) increases the likelihood of lighter eyes, in other words, the marker does not behave in an entirely dominant fashion. In fact, the effect of this marker is particularly strong for ΔE, indicating that heterozygous individuals are more likely to have central heterochromia. This observation has been reported in previous studies, where each copy of the ancestral allele (A) decreased the colour difference between the ciliary and pupillary zones (41). The same study also notes that rs12913832 may have both dominant/recessive and additive effects on iris colour variation, depending on which dimensions of colour space are being measured (41).

### Conclusion

A limitation of our study is the reliance on self-reported eye colour categories provided by participants in the CanPath cohorts. While practical, such categorical data may not reflect the full spectrum of iris pigmentation as accurately as quantitative measurement approaches (41,63,87). Additionally, participants may report their eye colour inaccurately or may not know which category to report (consider an individual with central heterochromia where the outer portion of their eye appears blue but there is a brown ring around the pupil). In this regard, we were fortunate to be able to replicate many of our signals in an independent European dataset for which quantitative iris colour measurements and a quantitative measurement of heterochromia were available. It is also important to note that the main goal of our project was to identify potential modifiers of the effect of rs12913832 polymorphism on eye colour, and not to evaluate the improvement on eye colour prediction based on the addition of these regions to current eye colour prediction panels, which ideally would require having access to additional large datasets with well-curated eye colour and genotype data. Finally, we fully acknowledge that solely focusing on individuals with European genetic ancestry limits the conclusions that can be drawn. Future studies would benefit from both increased size and diversity of study samples. Not only would it be interesting to see replication of results in other genetic ancestry groups, but an expansion of sample size and diversity may also draw attention to other genomic regions that are involved in the large spectrum of observed eye colour variation. In particular, recently admixed populations would provide an excellent opportunity to study in more detail gene-gene interactions shaping eye colour variation. The presence of uncharacterized alleles and epistatic effects may explain in part why forensic prediction tools for pigmentary traits have poorer performance in recently admixed populations than in European groups (88–90).

Together, these findings emphasize that the genetic modifiers of eye colour differ substantially between individuals with the rs12913832:GG and rs12913832:AA+AG backgrounds. When combined with expression information from GTEx, a clear pattern emerges from our GWAS data. For those with GG genotypes, who typically present with blue eyes, variants in core melanin synthesis genes (*TYR*, *TYRP1*), transporter proteins located on melanosomes (*SLC24A4*, *SLC45A2*), and potentially *TSPAN10* appear to contribute to darker-than-expected eye colour. Specifically, we observe that rs1126809:G (*TYR*), rs12896399:G (*SLC24A4*), and rs62075722:G (*TSPAN10*) all lead to increased expression of their respective genes (as reported in GTEx), and demonstrated eye colour darkening effects in GG stratified individuals. Conversely, among AA+AG individuals, lighter iris colours appear to result from a) other markers disrupting the expression or function of the *OCA2* gene and its product, independently of the *HERC2* rs12913832 polymorphism, and b) modulation by *IRF4*, which interacts with MITF and TFAP2α (transcription factor AP2α) to regulate *TYR* expression (91). For the two lead SNPs which had eQTLs in GTEx, we see that the rs12203592:T (*IRF4*) and rs1800407:T allele (*OCA2*) decrease expression in their respective genes, contributing to eye colour lightening in the AA+AG stratified individuals. Interestingly, *TYRP1* emerges in both genotype stratified groups. It seems that *TYRP1* may work both to counteract low expression *OCA2* levels in individuals with GG genotypes, and also decrease melanin levels in individuals with AA+AG genotypes.

In summary, this study advances our understanding of eye colour variability by identifying genetic variants that may modulate pigmentation in individuals with discordant *HERC2* rs12913832 genotypes. The identification of shared and background-specific loci underscores the polygenic and regulatory complexity of iris pigmentation. By integrating categorical self-report data with quantitative phenotype replication, our findings provide new insights into how moderating genetic factors contribute to subtle (and sometimes more pronounced) deviations from expected eye colour.

## Acknowledgements

The Canadian Partnership for Tomorrow’s Health (CanPath) research is only possible with the commitment of its research participants, its staff and its funders. The data used in this research were made available by CanPath with contributions from CARTaGENE, Alberta’s Tomorrow Project, the Ontario Health Study, the BC Generations Project, and Atlantic PATH. Computational analyses were carried out using the high-performance computing and storage resources of the Digital Research Alliance of Canada.

## Authors’ Roles

CLA: Formal Analysis, Writing – original draft, Writing – review and editing. FLD: Methodology, Writing – review and editing. ME: Data collection, Writing – review and editing. EJP: Conceptualization, Methodology, Writing – review and editing, Supervision.

## Competing Interests

N/A

## Funding

EJP received funding from the Natural Sciences and Engineering Research Council of Canada (***NSERC*** Discovery Grant).

## Data availability

The genotype information that support the findings of this study are available from CanPath but restrictions apply to the availability of these data, which were used under license for the current study, and so are not publicly available. Generated data are available from the corresponding author upon reasonable request. Scripts for the GWAS pipeline can be found at: https://github.com/cl-abba/GWAS-Pipeline/tree/main.

## Notes

### Competing Interest Statement

The authors have declared no competing interest.

### Summary of Updates

Data availability statement has been added to the manuscript.

