## Supplementary Material for "A comparative GWAS of eye colour in light and dark eye genetic backgrounds defined by *HERC2* rs12913832 polymorphism"

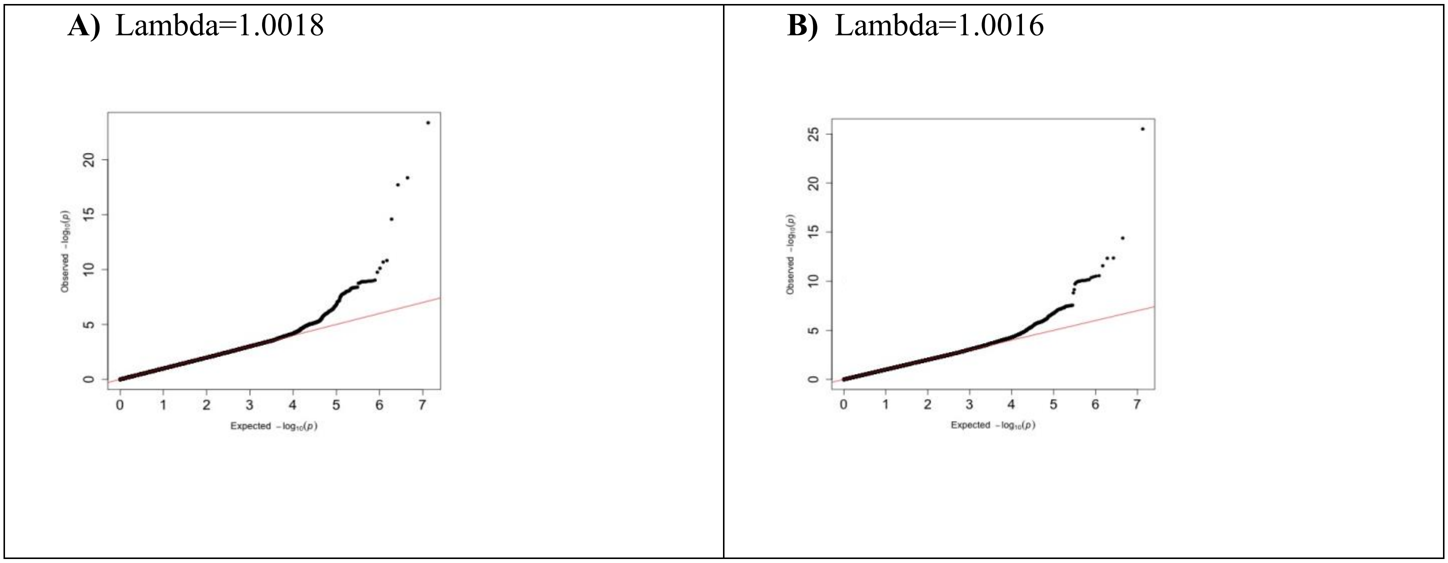

**Fig S1**. **QQ plots depicting observed vs expected P-values from meta-analyses conducted on rs12913832:AA+AG individuals.** **A)** Logistic model meta-analysis (UKBB+GSA arrays). **B)** Linear model meta-analysis (UKBB+GSA arrays).

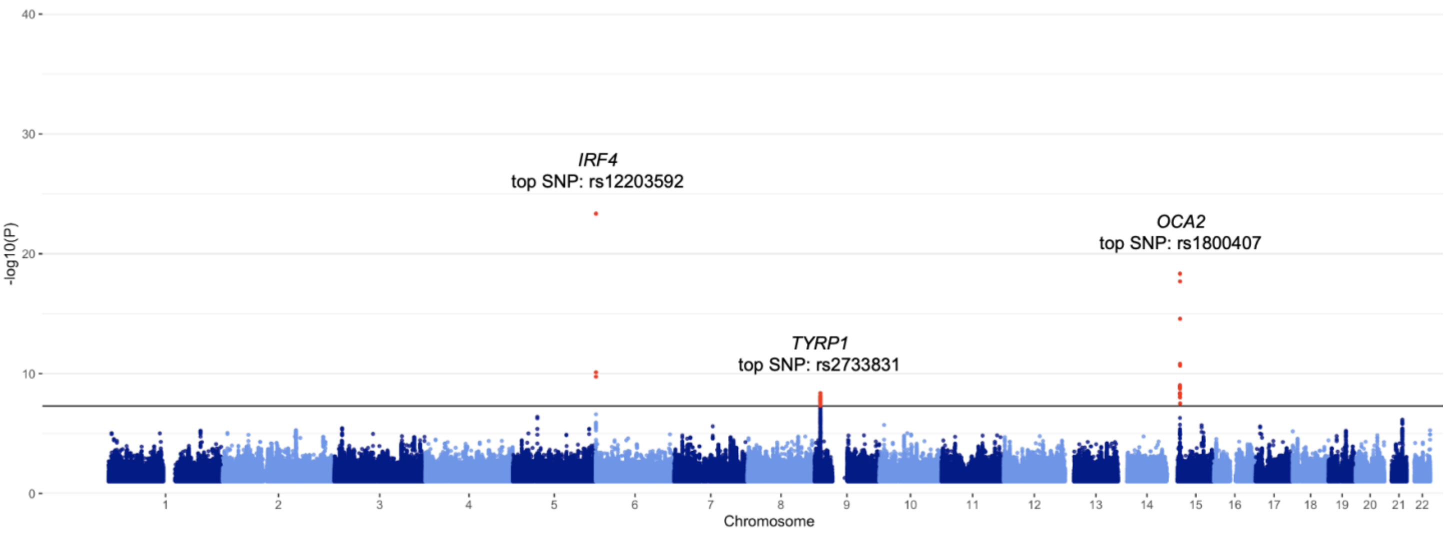

**Fig S2**. **Manhattan plot of eye color meta-analysis based on logistic model GWAS in rs12913832:AA+AG individuals (*N*=2,724) from the Canadian Partnership for Tomorrow's Health (CanPath) project genotyped with the Axiom UKBB and GSA arrays.** GWAS were performed with SAIGE (Zhou et al., 2018). Categorical eye colors coded as blue=1, hazel and brown=0. The meta-analysis was conducted with METASOFT (Zhao, n.d.). The black line indicates the genome-wide threshold (P<5E-8). Significant peaks are observed in chromosomes 6 (*IRF4*), 9 (*TYRP1*) and 15 (*OCA2*).

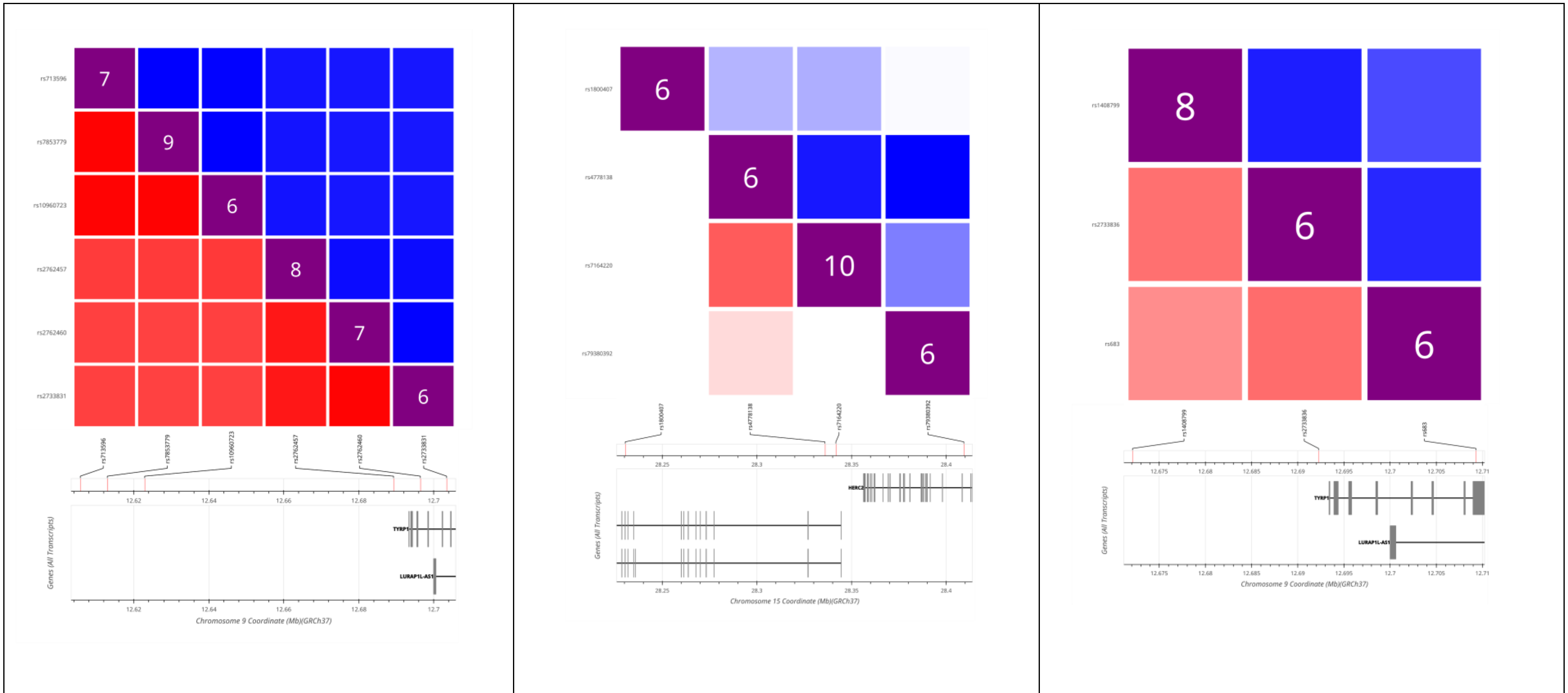

| A) ***TYRP1*** **(AA+AG background)** | B) ***HERC2/OCA2*** **(AA+AG background)** | C) ***TYRP1*** **(GG background, with prediction panel SNPs)** |
|---|---|---|

**Fig S3**. **LD matrices generated using the LDmatrix tool within LDlink (Machiela et al., 2015).** SNPs depicted represent those generated by FINEMAP (version 1.4) (Benner et al., 2016) that achieved a $\log_{10}$BF (Bayes Factor) greater than two in the **A**) *TYRP1* subset: rs7853779, rs10960723, rs2762457, rs2762460, rs2733831, **B**) HER2/OCA2 subset: rs1800407, rs7164220, rs4778138, rs79380392 and C) TYRP1 associated variant, rs2733836, with two SNPs previously associated with eye colour prediction, rs1408799 and rs683**.** The five EUR (European) populations were selected for reference (CEU [Utah Residents from North and West Europe], TSI [Toscani in Italia], FIN [Finnish in Finland], GBR [British in England and Scotland], and IBS [Iberian Population in Spain]). $R^2$ (red) is a measure of correlation of alleles for two genetic variants (ranges from 0 to 1, where a value of 0 indicates complete independence of alleles and a value of 1 indicates an allele of one variant that perfectly predicts an allele of another variant). Note: $R^2$ is sensitive to allele frequency. *D′* (blue) is an indicator of allelic segregation for two variants (ranges from 0 to 1, where a value of 0 indicates no linkage of alleles and a value of 1 indicates at least one haplotype combination that is not observed). A FORGEdb score is used for predicting which genetic variants are most likely to be regulatory, with the highest score of 9 computed from eQTL+ABC+TFmotif+CATO+DNase I hotspot+histone mark ChIP-seq. For more information regarding the calculations provided by LDlink, see https://ldlink.nci.nih.gov/?tab=help#Calculations. eQTL, expression Quantitative Trait Loci; LD, Linkage disequilibrium.

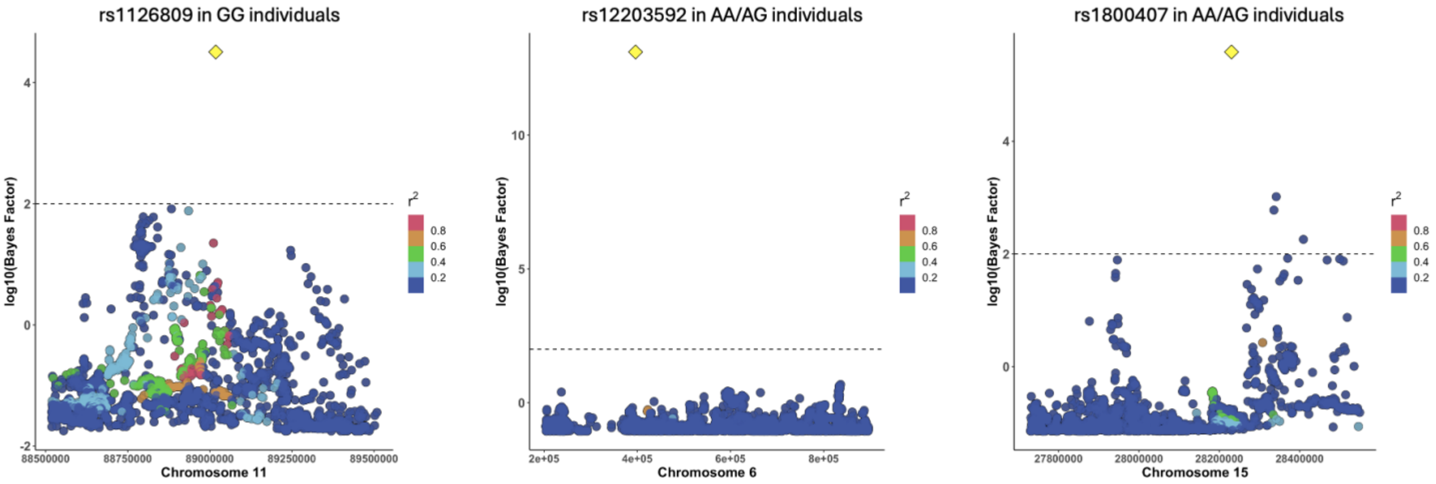

**Fig S4**. **Fine-map regional plots**. **A)** rs1126809 in GG individuals, **B)** rs1220352 in AA+AG individuals and rs1800407 in AA+AG individuals. Physical position along the chromosome is indicated on the x-axis and log10(Bayes Factor) is indicated on the y-axis. Lead SNP in each plot is highlighted with a yellow diamond. LD ($r^2$) correlations are shown in respect to the lead SNP. In each of these instances, the lead SNP is easily discernable among nearby SNPs, providing evidence for most likely causal SNP. Scripts for generated local plots can be found at https://github.com/cl-abba/GWAS-Pipeline/tree/main/Fine-mapping.

| Marker ID | Chr | Pos | Overlapped Gene | Type of Gene | Annotation | Nearest Upstream Gene | Nearest Downstream Gene | Roadmap Region Start | Roadmap Region End | Roadmap Feature Type Class | Roadmap Feature Type | Roadmap Epigenome | CADD PHRED Score | ClinVar Significance | ClinVar Phenotypes | Polyphen Prediction |
|---|---|---|---|---|---|---|---|---|---|---|---|---|---|---|---|---|
| **rs1126809** | chr11 | 89017961 | TYR | protein_coding | coding nonsyn,non-coding | None | None | | | | | | | Conflicting interpretations of pathogenicity | Albinism,Abnormality of metabolism/homeostasis,Elevated hepatic transaminases,Hypoplasia of the fovea,Slow decrease in visual acuity,Choroidal neovascularization,Oculocutaneous albinism,Autosomal recessive ocular albinism,Oculocutaneous albinism type 1B,Oculocutaneous albinism type 1, temperature sensitive,Skin/hair/eye pigmentation, variation in, 3,Skin/hair/eye pigmentation 3, blue/green eyes,Cutaneous malignant melanoma 8,Tyrosinase-negative oculocutaneous albinism,not specified,not provided | Probably damaging |
| **rs4904866** | chr14 | 92768503 | None | None | None | RNU6-366P | SLC24A4 | | | | | | 5.795 | | | |
| **rs12896399** | chr14 | 92773663 | None | None | None | RNU6-366P | SLC24A4 | | | | | | 0.3 | association | Skin/hair/eye pigmentation, variation in, 6 | |
| **rs12896471** | chr14 | 92773903 | None | None | None | RNU6-366P | SLC24A4 | | | | | | 1.045 | | | |
| **rs746586** | chr14 | 92775967 | None | None | None | RNU6-366P | SLC24A4 | | | | | | 1.781 | | | |
| **rs56137766** | chr14 | 92785941 | None | None | None | RNU6-366P | SLC24A4 | | | | | | 5.231 | | | |
| **rs35983729** | chr14 | 92787761 | None | None | None | RNU6-366P | SLC24A4 | | | | | | 5.697 | | | |
| **rs4144266** | chr14 | 92789205 | SLC24A4 | protein_coding | intronic,5upstream | None | None | | | | | | 7.619 | | | |
| **rs12883151** | chr14 | 92790077 | SLC24A4 | protein_coding | intronic,5upstream,5utr | None | None | | | | | | 8.981 | | | |
| **rs1800407** | chr15 | 28230318 | OCA2 | protein_coding | coding nonsyn | None | None | | | | | | | Benign/Likely benign | Oculocutaneous albinism,Skin/hair/eye pigmentation, variation in, 1,not specified | Probably damaging |
| **rs4778138** | chr15 | 28335820 | OCA2 | protein_coding | intronic | None | None | | | | | | 1.982 | | | |
| **rs7164220** | chr15 | 28341609 | OCA2 | protein_coding | intronic | None | None | 28340883 | 28342047 | Open Chromatin | DNase1 | foreskin melanocyte | 11.81 | | | |
| **rs79380392** | chr15 | 28409219 | HERC2 | protein_coding | intronic | None | None | | | | | | 5.332 | | | |
| **rs34635363** | chr17 | 79549250 | NPLOC4 | protein_coding | intronic | None | None | | | | | | 0.125 | | | |
| **rs62075722** | chr17 | 79611271 | TSPAN10 | polymorphic_pseudogene | non-coding intronic,intronic | None | None | | | | | | 6.286 | | | |
| **rs6420484** | chr17 | 79612397 | TSPAN10 | polymorphic_pseudogene | coding *nonsyn\|nonsyn,coding nonsyn\|nonsyn,non-coding | None | None | | | | | | | | | |
| **rs35407** | chr5 | 33946571 | SLC45A2 | protein_coding | 3downstream,intronic,3utr | None | None | | | | | | 2.185 | | | |

| | | | | | | | | | | | | | | | |
|---|---|---|---|---|---|---|---|---|---|---|---|---|---|---|---|
| **rs35395** | chr5 | 33948589 | SLC45A2 | protein_coding | intronic | None | None | | | | | | 7.193 | | |
| **rs16891982** | chr5 | 33951693 | SLC45A2 | protein_coding | 3utr,coding nonsyn\|nonsyn,coding *nonsyn\|nonsyn | None | None | | | | | | | Benign | Not specified |
| **rs35389** | chr5 | 33954880 | SLC45A2 | protein_coding | intronic | None | None | | | | | | 0.53 | | |
| **rs12203592** | chr6 | 396321 | IRF4 | protein_coding | non-coding intronic,intronic,3downstream | None | None | 394748 | 397443 | Open Chromatin | DNase1 | foreskin melanocyte | 11.78 | Affects | Skin/hair/eye pigmentation, variation in, 8 |
| **rs713596** | chr9 | 12605687 | None | None | None | RNU2-47P | TYRP1 | | | | | | 4.357 | | |
| **rs7853779** | chr9 | 12612893 | None | None | None | RNU2-47P | TYRP1 | | | | | | 7.814 | | |
| **rs10960723** | chr9 | 12622878 | None | None | None | RNU2-47P | TYRP1 | | | | | | 2.416 | | |
| **rs2762457** | chr9 | 12689313 | TYRP1 | protein_coding | intronic | None | None | | | | | | 1.391 | | |
| **rs2733836** | chr9 | 12692252 | TYRP1 | protein_coding | 5upstream,intronic | None | None | | | | | | 2.549 | | |
| **rs2762460** | chr9 | 12696478 | TYRP1 | protein_coding | 5upstream,intronic,3downstream | None | None | | | | | | 1.199 | | |
| **rs2733831** | chr9 | 12703484 | TYRP1 | protein_coding | non-coding intronic,intronic,3downstream | None | None | | | | | | 1.211 | | |

**Table S1**. **Summary of SNPnexus annotation output.** Lead SNPs highlighted.

**Marker ID**, reference SNP cluster ID (rsID);
**Chr**, chromosome;
**Pos**, physical position;
**Overlapped gene**, if the SNP directly overlaps with a gene, this column contains the name of the gene (HGNC system) to which the variant is overlapped;
**Overlapped gene type**, the type of gene (e.g., protein coding, miRNA, non coding, Pseudogene, snoRNA, lincRNA, etc.);
**Annotation**, summary of whether the variant overlapped with the coding, intronic or untranslated regions of the various transcript isoforms of the gene, as annotated from Ensembl gene system;
**Nearest upstream gene**, if variant is not overlapped with any gene, then the gene whose end position is nearest to the variant on the left (considering the alignment of genes on the positive strand as left-to-right);
**Nearest downstream gene**, if variant is not overlapped with any gene, then the gene whose start position is nearest to the variant on the right (considering the alignment of genes on the positive strand as left-to-right)
**Roadmap Region Start**, start position of the TFBS site in the chromosome;
**Roadmap Region End**, end position of the TFBS site in the chromosome;
**Roadmap Feature Type Class**, regulatory feature class;
**Roadmap Feature Type**, regulatory feature type;
**Roadmap Epigenome**, epigenome or cell name;
**CADD PHRED Score**, PHRED-like ($-10*\log_{10}$(rank/total)) scaled CADD-score ranking a variant relative to all possible substitutions of the human genome. A score≥10 indicates that it is predicted to be in the 10% most deleterious substitutions that you can do to the human genome, a score≥20 indicates the 1% most deleterious and so on.
**ClinVar Significance**, whether identified as Pathogenic or Benign or uncertain;
**ClinVar Phenotype**, List of phenotypes associated with the variant;
**PolyPhen Prediction**, PolyPhen predicted effect on protein based on the score. Possible values: Probably Damaging (score > 0.908), Possibly Damaging (0.446 < score <= 0.908), Benign (score <= 0.446).

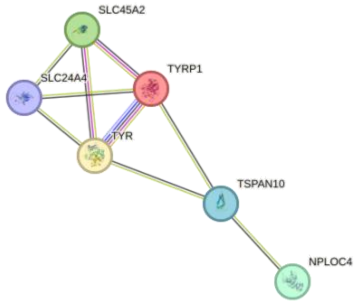

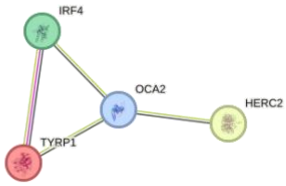

**(A) Gene network GG individuals**
Number of nodes=6
Number of edges=9
Average node degree=3
Average clustering coefficient = 0.778
Expected number of edges=0
PPI enrichment P-value=3.66E-15

**(B) Gene network AA+AG individuals**
Number of nodes=4
Number of edges=4
Average node degree=2
Average clustering coefficient = 0.833
Expected number of edges=0
PPI enrichment P-value=3.29E-6

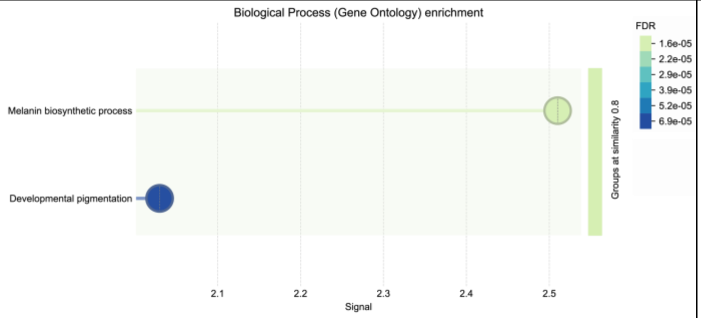

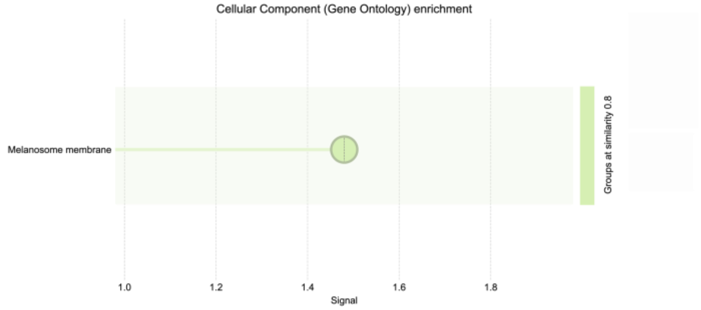

**(C) Biological process enrichment GG individuals**

**(D) Biological process enrichment AA+AG individuals**

**Fig S5**. **STRING (Szklarczyk et al., 2023) gene networks and enrichment ontology**. Network of genes derived from fine-mapped SNPs for each genotype-stratified GWAS (**A** and **B**) . Each coloured circle represents a gene. Genes which demonstrate known or predicted interactions are connected with a coloured edge: Aquamarine represents known interactions from curated databases. Magenta represents known interactions which have been experimentally determined. Dark green represents predicted interactions based on gene neighbourhood. Red represents predicted interactions based on gene fusions. Dark blue represents predicted interactions based on gene co-occurrence. Light green represents connections made via textmining. Black represents connections made via co-expression. Light blue represents connections made via protein homology. **C** and **D** illustrate the enrichment analysis provided by STRING. The x-axis represents the enrichment signal. The signal is defined as a weighted harmonic mean between the observed/expected ratio and -log(FDR). FDR tends to emphasize larger terms due to their potential for achieving lower p-values, while the observed/expected ratio highlights smaller terms, which have a high foreground to background ratio but cannot achieve low FDR values due to their size. The signal measure seeks to balance both metrics for more intuitive ordering of enriched terms. The color gradient corresponds to FDR values, with darker shades representing more significant enrichment. The size of each bubble represents the number of genes contributing to the enrichment of a particular process. The green background highlights the most significantly enriched processes, while the blue region represents terms with relatively lower significance.

| #category | term ID | term description | observed gene count | background gene count | strengt h | signa l | false discovery rate | matching proteins in your network (labels) |
|---|---|---|---|---|---|---|---|---|
| **GG individuals** | | | | | | | | |
| GO Process | GO:0042438 | Melanin biosynthetic process | 3 | 14 | 2.85 | 2.51 | 0.00017 | TYR,SLC45A2,TYRP1 |
| GO Process | GO:0048066 | Developmental pigmentation | 3 | 44 | 2.35 | 2.03 | 0.00069 | TYR,SLC45A2,TYRP1 |
| GO Function | GO:0004503 | Tyrosinase activity | 2 | 2 | 3.52 | 1.79 | 0.0023 | TYR,TYRP1 |
| GO Component | GO:0033162 | Melanosome membrane | 3 | 18 | 2.74 | 2.88 | 4.26E-05 | TYR,SLC45A2,TYRP1 |
| STRING clusters | CL:14861 | Melanin biosynthetic process, and Domain of unknown function (DUF3371) | 3 | 13 | 2.88 | 2.92 | 4.01E-05 | TYR,SLC45A2,TYRP1 |
| STRING clusters | CL:14863 | Oculocutaneous albinism type IV, and Agouti | 2 | 5 | 3.12 | 1.76 | 0.0025 | SLC45A2,TYRP1 |
| KEGG | hsa00350 | Tyrosine metabolism | 2 | 35 | 2.27 | 1.16 | 0.0172 | TYR,TYRP1 |
| Reactome | HSA-5662702 | Melanin biosynthesis | 3 | 5 | 3.29 | 3.84 | 2.00E-06 | TYR,SLC45A2,TYRP1 |
| WikiPathways | WP4941 | GPR143 in melanocytes and retinal pigment epithelium cells | 2 | 29 | 2.35 | 1.03 | 0.028 | TYR,TYRP1 |
| Monarch | EFO:0003924 | Hair color | 6 | 120 | 2.22 | 4.98 | 7.40E-10 | TYR,SLC45A2,NPLOC4,TYRP1,SLC24A4,TSPAN1 0 |
| Monarch | EFO:0003949 | Eye color | 5 | 35 | 2.67 | 5.73 | 7.40E-10 | TYR,SLC45A2,NPLOC4,SLC24A4,TSPAN10 |
| Monarch | EFO:0007822 | Hair colour measurement | 5 | 249 | 1.82 | 2.62 | 6.20E-06 | TYR,SLC45A2,NPLOC4,SLC24A4,TSPAN10 |
| Monarch | HP:0000635 | Blue irides | 3 | 19 | 2.71 | 2.83 | 5.08E-05 | TYR,SLC45A2,TYRP1 |
| Monarch | HP:0001022 | Albinism | 3 | 18 | 2.74 | 2.83 | 5.08E-05 | TYR,SLC45A2,TYRP1 |
| Monarch | HP:0011358 | Generalized hypopigmentation of hair | 3 | 41 | 2.38 | 2.27 | 0.00028 | TYR,SLC45A2,TYRP1 |
| Monarch | HP:0007730 | Iris hypopigmentation | 3 | 48 | 2.31 | 2.16 | 0.0004 | TYR,SLC45A2,TYRP1 |
| Monarch | EFO:0003784 | Skin pigmentation | 3 | 82 | 2.08 | 1.71 | 0.0017 | TYR,SLC45A2,SLC24A4 |
| Monarch | HP:0025551 | Optic nerve misrouting | 2 | 5 | 3.12 | 1.87 | 0.0017 | SLC45A2,TYRP1 |
| Monarch | HP:0200098 | Absent skin pigmentation | 2 | 6 | 3.04 | 1.84 | 0.0019 | TYR,TYRP1 |
| Monarch | HP:0003764 | Nevus | 3 | 109 | 1.96 | 1.57 | 0.0025 | TYR,SLC45A2,TYRP1 |
| Monarch | HP:0002297 | Red hair | 2 | 9 | 2.86 | 1.71 | 0.0029 | TYR,TYRP1 |
| Monarch | EFO:0010554 | Retinal vasculature measurement | 2 | 10 | 2.82 | 1.67 | 0.0033 | NPLOC4,TSPAN10 |

| | | | | | | | | |
|---|---|---|---|---|---|---|---|---|
| Monarch | HP:0002226 | White eyebrow | 2 | 11 | 2.78 | 1.63 | 0.0037 | TYR,TYRP1 |
| Monarch | HP:0002227 | White eyelashes | 2 | 13 | 2.7 | 1.56 | 0.0047 | TYR,TYRP1 |
| Monarch | HP:0001107 | Ocular albinism | 2 | 15 | 2.64 | 1.49 | 0.0058 | TYR,SLC45A2 |
| Monarch | EFO:0003963 | Freckles | 2 | 18 | 2.56 | 1.42 | 0.0074 | TYR,SLC45A2 |
| Monarch | HP:0011364 | White hair | 2 | 20 | 2.52 | 1.37 | 0.0087 | TYR,SLC45A2 |
| Monarch | EFO:0600004 | Retinal nerve fibre layer thickness measurement | 2 | 21 | 2.5 | 1.35 | 0.0091 | TYR,NPLOC4 |
| Monarch | EFO:0600005 | Ganglion cell inner plexiform layer thickness measurement | 2 | 21 | 2.5 | 1.35 | 0.0091 | TYR,NPLOC4 |
| Monarch | EFO:0004695 | Intraocular pressure measurement | 3 | 211 | 1.67 | 1.13 | 0.0107 | TYR,NPLOC4,TSPAN10 |
| Monarch | HP:0007894 | Hypopigmentation of the fundus | 2 | 24 | 2.44 | 1.3 | 0.0107 | TYR,SLC45A2 |
| Monarch | EFO:0007906 | Synophrys measurement | 2 | 26 | 2.4 | 1.27 | 0.0118 | SLC45A2,SLC24A4 |
| Monarch | EFO:0004731 | Eye measurement | 4 | 804 | 1.21 | 0.84 | 0.014 | TYR,SLC45A2,NPLOC4,TSPAN10 |
| Monarch | EFO:0009464 | Corneal disease | 2 | 29 | 2.35 | 1.22 | 0.014 | NPLOC4,TSPAN10 |
| Monarch | EFO:1002040 | Corneal astigmatism | 2 | 29 | 2.35 | 1.22 | 0.014 | NPLOC4,TSPAN10 |
| Monarch | HP:0000483 | Astigmatism | 3 | 239 | 1.62 | 1.05 | 0.014 | TYR,NPLOC4,TSPAN10 |
| Monarch | HP:0007750 | Hypoplasia of the fovea | 2 | 29 | 2.35 | 1.22 | 0.014 | TYR,SLC45A2 |
| Monarch | HP:0008060 | Aplasia/Hypoplasia of the fovea | 2 | 29 | 2.35 | 1.22 | 0.014 | TYR,SLC45A2 |
| Monarch | EFO:0007009 | Skin pigmentation measurement | 2 | 32 | 2.31 | 1.21 | 0.0142 | TYR,SLC45A2 |
| Monarch | HP:0001480 | Freckling | 2 | 38 | 2.24 | 1.13 | 0.0188 | TYR,TYRP1 |
| Monarch | EFO:0009180 | Rosacea severity measurement | 2 | 47 | 2.15 | 1.03 | 0.0256 | SLC45A2,SLC24A4 |
| Monarch | EFO:0010176 | Keratinocyte carcinoma | 2 | 47 | 2.15 | 1.03 | 0.0256 | TYR,SLC45A2 |
| Monarch | EFO:0004279 | Suntan | 2 | 49 | 2.13 | 1.02 | 0.0265 | TYR,SLC45A2 |
| Monarch | EFO:0004632 | Nevus count | 2 | 57 | 2.06 | 0.95 | 0.0341 | TYR,SLC45A2 |
| Monarch | HP:0004328 | Abnormal anterior eye segment morphology | 4 | 1106 | 1.07 | 0.64 | 0.0348 | TYR,SLC45A2,NPLOC4,TYRP1 |
| Monarch | HP:0012373 | Abnormal eye physiology | 5 | 2405 | 0.83 | 0.5 | 0.0366 | TYR,SLC45A2,NPLOC4,TYRP1,TSPAN10 |
| Monarch | EFO:0009764 | Eye colour measurement | 2 | 66 | 2 | 0.88 | 0.0426 | TYR,SLC45A2 |
| DISEASES | DOID:0070097 | Oculocutaneous albinism type III | 3 | 6 | 3.22 | 3.51 | 6.05E-06 | TYR,SLC45A2,TYRP1 |

| | | | | | | | | |
|---|---|---|---|---|---|---|---|---|
| DISEASES | DOID:0070098 | Oculocutaneous albinism type IV | 3 | 6 | 3.22 | 3.51 | 6.05E-06 | TYR,SLC45A2,TYRP1 |
| DISEASES | DOID:1909 | Melanoma | 3 | 46 | 2.33 | 2.21 | 0.00033 | TYR,SLC45A2,TYRP1 |
| DISEASES | DOID:0070095 | Oculocutaneous albinism type IB | 2 | 5 | 3.12 | 1.91 | 0.0015 | TYR,SLC45A2 |
| DISEASES | DOID:8923 | Skin melanoma | 2 | 12 | 2.74 | 1.52 | 0.0054 | TYR,SLC45A2 |
| TISSUES | BTO:0000849 | Melanoma cell line | 4 | 180 | 1.86 | 2 | 0.00026 | TYR,SLC45A2,NPLOC4,TYRP1 |
| COMPARTMENTS | GOCC:0033162 | Melanosome membrane | 3 | 14 | 2.85 | 3.06 | 2.45E-05 | TYR,SLC45A2,TYRP1 |
| UniProt Keywords | KW-0470 | Melanin biosynthesis | 3 | 7 | 3.15 | 3.95 | 1.26E-06 | TYR,SLC45A2,TYRP1 |
| UniProt Keywords | KW-0015 | Albinism | 3 | 19 | 2.71 | 3.33 | 8.10E-06 | TYR,SLC45A2,TYRP1 |
| UniProt Keywords | KW-0186 | Copper | 2 | 65 | 2 | 0.91 | 0.0379 | TYR,TYRP1 |
| UniProt Keywords | KW-0503 | Monooxygenase | 2 | 79 | 1.92 | 0.88 | 0.0416 | TYR,TYRP1 |
| Pfam | PF00264 | Common central domain of tyrosinase | 2 | 3 | 3.34 | 1.86 | 0.0018 | TYR,TYRP1 |
| InterPro | IPR002227 | Tyrosinase copper-binding domain | 2 | 3 | 3.34 | 1.54 | 0.0055 | TYR,TYRP1 |
| InterPro | IPR008922 | Di-copper centre-containing domain superfamily | 2 | 3 | 3.34 | 1.54 | 0.0055 | TYR,TYRP1 |
| **AA_AG individuals** | | | | | | | | |
| GO Component | GO:0033162 | Melanosome membrane | 2 | 18 | 2.74 | 1.48 | 0.012 | OCA2,TYRP1 |
| Reactome | HSA-5662702 | Melanin biosynthesis | 2 | 5 | 3.29 | 2.2 | 0.0015 | OCA2,TYRP1 |
| WikiPathways | WP3998 | Prader-Willi and Angelman syndrome | 2 | 62 | 2.2 | 0.99 | 0.0485 | HERC2,OCA2 |
| Monarch | EFO:0003924 | Hair color | 4 | 120 | 2.22 | 3.21 | 1.82E-05 | HERC2,OCA2,IRF4,TYRP1 |
| Monarch | HP:0000635 | Blue irides | 3 | 19 | 2.89 | 3.57 | 1.96E-05 | HERC2,OCA2,TYRP1 |
| Monarch | EFO:0007009 | Skin pigmentation measurement | 3 | 32 | 2.66 | 3.14 | 6.26E-05 | HERC2,OCA2,IRF4 |
| Monarch | EFO:0003949 | Eye color | 3 | 35 | 2.63 | 3.12 | 6.45E-05 | HERC2,OCA2,IRF4 |
| Monarch | EFO:0004279 | Suntan | 3 | 49 | 2.48 | 2.82 | 0.00014 | HERC2,OCA2,IRF4 |
| Monarch | EFO:0009764 | Eye colour measurement | 3 | 66 | 2.35 | 2.55 | 0.00029 | HERC2,OCA2,IRF4 |
| Monarch | HP:0025551 | Optic nerve misrouting | 2 | 5 | 3.29 | 2.41 | 0.00079 | OCA2,TYRP1 |
| Monarch | HP:0200098 | Absent skin pigmentation | 2 | 6 | 3.22 | 2.34 | 0.00096 | OCA2,TYRP1 |
| Monarch | HP:0002297 | Red hair | 2 | 9 | 3.04 | 2.14 | 0.0017 | OCA2,TYRP1 |
| Monarch | HP:0002226 | White eyebrow | 2 | 11 | 2.95 | 2.04 | 0.0023 | OCA2,TYRP1 |

| | | | | | | | | |
|---|---|---|---|---|---|---|---|---|
| Monarch | HP:0002227 | White eyelashes | 2 | 13 | 2.88 | 1.97 | 0.0028 | OCA2,TYRP1 |
| Monarch | HP:0001022 | Albinism | 2 | 18 | 2.74 | 1.78 | 0.0048 | OCA2,TYRP1 |
| Monarch | EFO:0007822 | Hair colour measurement | 3 | 249 | 1.77 | 1.44 | 0.0062 | HERC2,OCA2,IRF4 |
| Monarch | EFO:0007906 | Synophrys measurement | 2 | 26 | 2.58 | 1.58 | 0.0084 | HERC2,IRF4 |
| Monarch | HP:0001480 | Freckling | 2 | 38 | 2.41 | 1.38 | 0.0147 | OCA2,TYRP1 |
| Monarch | HP:0011358 | Generalized hypopigmentation of hair | 2 | 41 | 2.38 | 1.35 | 0.0162 | OCA2,TYRP1 |
| Monarch | EFO:0005677 | Puberty onset measurement | 2 | 47 | 2.32 | 1.27 | 0.0201 | HERC2,IRF4 |
| Monarch | EFO:0009180 | Rosacea severity measurement | 2 | 47 | 2.32 | 1.27 | 0.0201 | HERC2,IRF4 |
| Monarch | EFO:0010176 | Keratinocyte carcinoma | 2 | 47 | 2.32 | 1.27 | 0.0201 | HERC2,IRF4 |
| Monarch | HP:0007730 | Iris hypopigmentation | 2 | 48 | 2.31 | 1.27 | 0.0201 | OCA2,TYRP1 |
| Monarch | EFO:0004632 | Nevus count | 2 | 57 | 2.24 | 1.21 | 0.0238 | HERC2,IRF4 |
| DISEASES | DOID:1909 | Melanoma | 4 | 46 | 2.63 | 4.89 | 1.69E-07 | HERC2,OCA2,IRF4,TYRP1 |
| DISEASES | DOID:0070097 | Oculocutaneous albinism type III | 2 | 6 | 3.22 | 2.24 | 0.0013 | OCA2,TYRP1 |
| DISEASES | DOID:0070098 | Oculocutaneous albinism type IV | 2 | 6 | 3.22 | 2.24 | 0.0013 | OCA2,TYRP1 |
| COMPARTMENTS | GOCC:0033162 | Melanosome membrane | 2 | 14 | 2.85 | 1.6 | 0.0085 | OCA2,TYRP1 |
| UniProt Keywords | KW-0015 | Albinism | 2 | 19 | 2.71 | 1.8 | 0.0044 | OCA2,TYRP1 |

**Table S2**. **Summary of functional enrichments in gene networks for GG and AA+AG genetic backgrounds.** Enrichment table columns (explanations are as provided on STRING (Szklarczyk et al., 2023)):

**Count In Network:**The first number indicates how many proteins in your network are annotated with a particular term. The second number indicates how many proteins in total (in your network and in the background) have this term assigned. You can click on the numbers to see the network view of the gene sets behind them.

**Strength:** Log10(observed / expected). This measure describes how large the enrichment effect is. It's the ratio between i) the number of proteins in your network that are annotated with a term and ii) the number of proteins that we expect to be annotated with this term in a random network of the same size.

**Signal:** The signal is defined as a weighted harmonic mean between the observed/expected ratio and -log(FDR). FDR tends to emphasize larger terms due to their potential for achieving lower p-values, while the observed/expected ratio highlights smaller terms, which have a high foreground to background ratio but cannot achieve low FDR values due to their size. The signal measure seeks to balance both metrics for more intuitive ordering of enriched terms.

**False Discovery Rate:** This measure describes how significant the enrichment is. Shown are p-values corrected for multiple testing within each category using the Benjamini–Hochberg procedure.

| Quantitative measure | Chr | Pos | ID | A1 | A2 | A1 frequency | B | SE | P |
|---|---|---|---|---|---|---|---|---|---|
| L (Lightness dimension: 0 (black) to 100 (white)) | **GG** | | | | | | | | |
| | 5 | 33954880 | rs35389 | G | A | 0.0393836 | 0.958527 | 1.04402 | 0.359339 |
| | 9 | 12692252 | rs2733836 | T | G | 0.460616 | 0.00876916 | 0.422575 | 0.983458 |
| | 11 | 89017961 | rs1126809 | A | G | 0.270548 | 1.43427 | 0.456852 | 0.00187087 |
| | 14 | 92773663 | rs12896399 | T | G | 0.424658 | 0.068321 | 0.40571 | 0.86639 |
| | 17 | 79611271 | rs62075722 | A | G | 0.354452 | 0.550985 | 0.411158 | 0.18129 |
| | **AA+AG** | | | | | | | | |
| | 6 | 396321 | rs12203592 | T | C | 0.155642 | 2.45411 | 0.754708 | 0.001306 |
| | 9 | 12612893 | rs7853779 | A | G | 0.420233 | -0.4656 | 0.585478 | 0.427223 |
| | 15 | 28230318 | rs1800407 | T | C | 0.105058 | 3.49465 | 0.938079 | 0.000241 |
| | 15 | 28335820 | rs4778138 | G | A | 0.264591 | -2.0395 | 0.67052 | 0.002604 |
| | 15 | 28365618 | rs12913832 | G | A | 0.412451 | 5.25802 | 0.997904 | 2.97E-07 |
| a (Green [-] to red [+]) | **GG** | | | | | | | | |
| | 5 | 33954880 | rs35389 | G | A | 0.0393836 | -0.186845 | 0.1379 | 0.176493 |
| | 9 | 12692252 | rs2733836 | T | G | 0.460616 | -0.00602619 | 0.0564623 | 0.915077 |
| | 11 | 89017961 | rs1126809 | A | G | 0.270548 | -0.120172 | 0.0613105 | 0.0509459 |
| | 14 | 92773663 | rs12896399 | T | G | 0.424658 | -0.0581953 | 0.0547672 | 0.28885 |
| | 17 | 79611271 | rs62075722 | A | G | 0.354452 | 0.0688326 | 0.0553151 | 0.214367 |
| | **AA+AG** | | | | | | | | |
| | 6 | 396321 | rs12203592 | T | C | 0.155642 | -0.38727 | 0.089875 | 2.34E-05 |
| | 9 | 12612893 | rs7853779 | A | G | 0.420233 | 0.024898 | 0.0715 | 0.727964 |
| | 15 | 28230318 | rs1800407 | T | C | 0.105058 | -0.31536 | 0.115791 | 0.006905 |
| | 15 | 28335820 | rs4778138 | G | A | 0.264591 | 0.06769 | 0.081711 | 0.408211 |
| | 15 | 28365618 | rs12913832 | G | A | 0.412451 | -0.37672 | 0.126653 | 0.003217 |
| b (Blue [-] to yellow [+]) | **GG** | | | | | | | | |
| | 5 | 33954880 | rs35389 | G | A | 0.0393836 | 0.0650803 | 0.142307 | 0.647781 |
| | 9 | 12692252 | rs2733836 | T | G | 0.460616 | 0.0360806 | 0.0580668 | 0.534848 |
| | 11 | 89017961 | rs1126809 | A | G | 0.270548 | -0.251696 | 0.0617662 | 5.95E-05 |
| | 14 | 92773663 | rs12896399 | T | G | 0.424658 | -0.22453 | 0.0549086 | 5.61E-05 |
| | 17 | 79611271 | rs62075722 | A | G | 0.354452 | 0.203995 | 0.0558043 | 0.000304612 |
| | **AA+AG** | | | | | | | | |
| | 6 | 396321 | rs12203592 | T | C | 0.155642 | 0.251521 | 0.0846127 | 0.00323522 |
| | 9 | 12612893 | rs7853779 | A | G | 0.420233 | -0.0427731 | 0.0660667 | 0.51794 |
| | 15 | 28230318 | rs1800407 | T | C | 0.105058 | -0.15877 | 0.108144 | 0.1433 |
| | 15 | 28335820 | rs4778138 | G | A | 0.264591 | -0.0433989 | 0.0755982 | 0.566424 |
| | 15 | 28365618 | rs12913832 | G | A | 0.412451 | -0.0215752 | -0.0215752 | 0.856396 |
| ΔE (heterochromia measure) | **GG** | | | | | | | | |
| | 5 | 33954880 | rs35389 | G | A | 0.0393836 | 0.243538 | 0.191137 | 0.203628 |
| | 9 | 12692252 | rs2733836 | T | G | 0.460616 | 0.0913852 | 0.0780487 | 0.242611 |
| | 11 | 89017961 | rs1126809 | A | G | 0.270548 | 0.0159388 | 0.0855046 | 0.852254 |
| | 14 | 92773663 | rs12896399 | T | G | 0.424658 | -0.126292 | 0.0756678 | 0.0961895 |
| | 17 | 79611271 | rs62075722 | A | G | 0.354452 | 0.119658 | 0.0765244 | 0.118987 |
| | **AA+AG** | | | | | | | | |
| | 6 | 396321 | rs12203592 | T | C | 0.155642 | 0.377876 | 0.120246 | 0.00187289 |

| | | | | | | | | | |
|---|---|---|---|---|---|---|---|---|---|
| | 9 | 12612893 | rs7853779 | A | G | 0.420233 | -0.0177142 | 0.0941452 | 0.850903 |
| | 15 | 28230318 | rs1800407 | T | C | 0.105058 | 0.247194 | 0.153862 | 0.109382 |
| | 15 | 28335820 | rs4778138 | G | A | 0.264591 | -0.177423 | 0.107142 | 0.098959 |
| | 15 | 28365618 | rs12913832 | G | A | 0.412451 | 0.848141 | 0.161075 | 0.000000297 |
| | **GG** | | | | | | | | |
| | 5 | 33954880 | rs35389 | G | A | 0.0393836 | 0.13956 | 0.0940053 | 0.138761 |
| | 9 | 12692252 | rs2733836 | T | G | 0.460616 | 0.01671 | 0.0381273 | 0.661526 |
| | 11 | 89017961 | rs1126809 | A | G | 0.270548 | -0.0999551 | 0.0415218 | 0.0167093 |
| | 14 | 92773663 | rs12896399 | T | G | 0.424658 | -0.05485 | 0.0364749 | 0.133751 |
| SREC1 (linear model) | 17 | 79611271 | rs62075722 | A | G | 0.354452 | -0.0393929 | 0.0371534 | 0.289919 |
| | **AA+AG** | | | | | | | | |
| | 6 | 396321 | rs12203592 | T | C | 0.155642 | -0.41453 | 0.088375 | 4.49E-06 |
| | 9 | 12612893 | rs7853779 | A | G | 0.420233 | 0.060526 | 0.070035 | 0.388294 |
| | 15 | 28230318 | rs1800407 | T | C | 0.105058 | -0.39477 | 0.112578 | 0.000538 |
| | 15 | 28335820 | rs4778138 | G | A | 0.264591 | 0.230776 | 0.080384 | 0.004445 |
| | 15 | 28365618 | rs12913832 | G | A | 0.412451 | -0.49868 | 0.121846 | 5.76E-05 |

**Table S3. Replication results in independent European sample using quantitative eye colour measures**. Eye colour was quantitatively assessed in 549 individuals of European ancestry using high-resolution images analyzed in CIELAB color space (Edwards et al., 2015). The L value indicates lightness (0 = black, 100 = white), while a and b* represent green–red and blue–yellow variation, respectively. Blue eyes are typically characterized by higher L*, negative a*, and negative b* values. The amount of central heterochromia (i.e. differences in iris colour between the inner and outer portions of the iris) was measured by a colour metric (ΔE) that represents differences in the average colour of the ciliary zone (outer iris region) and the average colour of the pupillary zone Inner iris region). Eye colours were categorized as: 0=Light blue, Light gray or light green; 1=Blue, gray or green, 2=Hazel or light brown, 3=Dark brown, 4=Brownish black. Markers which illustrated nominally significant P-values are highlighted in red, and markers which survive Bonferroni correction (0.01 in GG stratified and 0.0125 in AA+AG stratified) are highlighted in green.

**Chr**, chromosome;
**Pos**, Physical position;
**ID**, Marker ID (rsID);
**A1**, effect allele: the allele whose effects in relation to eye color are being studied;
**A2**, alternative allele;
**A1 frequency**, fraction of chromosomes in population that carry the effect allele;
**Beta**, per unit increase (if positive) or decrease (if negative) in the outcome
**SE**, standard error;
**P**, parameter of statistical significance used to determine the certainty of an association.

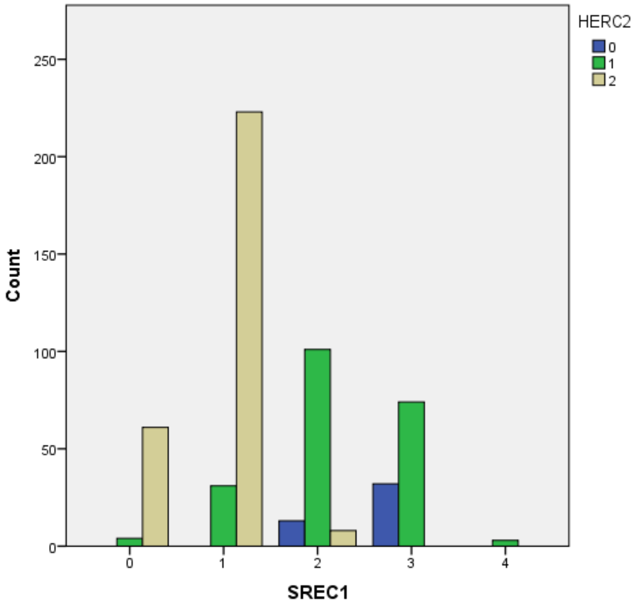

**Fig S6. Distributions of rs12913832 (*HERC2*) genotype across eye colour categories in the linear model.** Referring to the legend, a *HERC2* value of 0 represents the AA genotype, 1 represents a heterozygous genotype, and 2 represents the GG genotype. Eye colour categories are 0=Light blue, Light gray or light green; 1=Blue, gray or green, 2=Hazel or light brown, 3=Dark brown, 4=Brownish black).
